# Genetic and environmental influence on the human functional connectome

**DOI:** 10.1101/277996

**Authors:** Andrew E. Reineberg, Alexander S. Hatoum, John K. Hewitt, Marie T. Banich, Naomi P. Friedman

## Abstract

Detailed mapping of genetic and environmental influences on the functional connectome is a crucial step toward developing intermediate phenotypes between genes and clinical diagnoses or cognitive abilities. We analyze resting-state data from two, adult twin samples - 390 twins from the Colorado Longitudinal Twin Sample and 422 twins from the Human Connectome Project - to examine genetic and environmental influence on all pairwise functional connections between 264 brain regions (~35,000 functional connections). Non-shared environmental influence was high, genetic influence was moderate, and shared environmental influence was weak-to-moderate across the connectome. The brain’s genetic organization is diverse and not as one would expect based solely on structure evident in non-genetically informative data or lower-resolution data. As follow-up, we make novel classifications of functional connections and examine highly-localized connections with particularly strong genetic influence. This high-resolution genetic taxonomy of brain connectivity will be useful in understanding genetic influences on brain disorders.

---

The functional connectome refers to intrinsically correlated activity between brain regions when individuals are not engaged in a particular task (i.e., measured during the “resting state”; [Fox and Raichle, 2007]). Patterns within the functional connectome are associated with clinical diagnoses (for reviews see [Greicius, 2008, Zhang and Raichle, 2010]) and individual differences in cognitive abilities (for a broad review of 125 studies see [Vaidya and Gordon, 2013]). Recent work has showcased reliable and generalizable predictive models of individual differences in behavior that utilize many measurements of the connectome as features [Rosenberg et al., 2016, Finn et al., 2015]. These patterns of connectivity may be candidate intermediate phenotypes between genes and traits (i.e., endophenotypes; [Hall and Smoller, 2010, Kendler and Neale, 2010]) if genetic influences exist. However, prior work has only quantified genetic and environmental influences on the connectome at the level of large regions/networks of interest. In the current study, we perform a highly detailed analysis of the etiology of functional brain connections and reveal a high degree of diversity in genetic influence across the connectome.

Genetic analyses of functional connections could span several units of analysis - from connections between a small number of large networks with correlated activity [Yeo et al., 2011] and related function [Smith et al., 2009] to connections between nearly a million individual voxels. Due to the computational power needed to perform classic twin models at the level of voxels or small regions, current efforts have focused on quantifying genetic and environmental influence on global summary measures of functional connectivity, resting-state networks (i.e., large and spatially-separated groups of regions that are all moderately correlated at rest and thus appropriate to model as a single unit), or large regions of interest (ROIs). At the coarsest level of detail, several studies have revealed moderate heritability (i.e., “h^2^” or the proportion of phenotypic variance explained by genetic variance; h^2^ = 0.43 - 0.64) of the degree to which an individual’s connectome is globally efficient (i.e., maximizes information transfer while reducing long path lengths and unnecessary connections; [van den Heuvel et al., 2013, Fornito et al., 2011, Sinclair et al., 2015]). However, while global efficiency may be an informative phenotype, it may not be a thorough summary of the entire connectome, possibly summarizing only connections amongst the brains’ densely connected and metabolically costly hub regions [Heuvel et al., 2012].

At the level of networks, there was substantial early interest in the genetics of the default network, a set of regions implicated in internal mentation functions [Andrews-Hanna, 2011], perhaps due to its involvement in a variety of clinical disorders such as schizophrenia, depression, and attention deficit hyperactivity disorder [Whitfield-Gabrieli and Ford, 2012, Anticevic et al., 2012, Kaiser et al., 2016, Mattfeld et al., 2014, Whitfield-Gabrieli et al., 2009]. The default network is moderately heritable as a whole (h^2^ = 0.42), while connectivity of subcomponents of the default network are weakly-to-moderately heritable (h^2^ = 0.10 - 0.42; [Glahn et al., 2010]). Other work has reported moderate heritability of a precuneus-dorsal posterior cingulate network, visual network, default network, frontoparietal network, and dorsal attention network (h^2^ = 0.23 - 0.65), non-significant heritability for the salience and sensory-somatomotor networks, and evidence of environmental effects on functional connectivity between all networks [Yang et al., 2016]. Finally, a recent study investigated the genetic etiology of functional connections among seven networks and pairwise connections between 51 brain areas, finding moderate-to-strong heritability of seven networks (h^2^ = ~0.60 - ~0.75) using a linear mixed effects model approach to account for unreliability across multiple resting-state scans [Ge et al., 2017]. At the level of the 51-region parcellation, the authors found heritability estimates for components of some network, such as the default network, were consistent, but also found evidence of heterogeneity for regions of other networks such as the limbic and cognitive control networks. In summary, existing studies have provided heritability estimates for functional connectivity at global, network, or large ROI levels of analysis.

Although coarser levels of analysis are undoubtedly informative, they are not without caveats. First, large networks have only vague overarching functional labels (such as “vision”) as opposed to distinct functional labels ascribed to regions of finely detailed parcellations (e.g., those that contain 200 - 500 regions). Second, the anatomical literature indicates heritability may be overestimated for larger versus smaller pieces of cortex [Eyler et al., 2012]. Third, examining heritability at the network level assumes that areas within the networks are homogeneous in terms of their genetic connections to areas in other networks. Fourth, individual differences in within-network connectivity cannot be examined, and these individual differences may have important implications for behavior (i.e., as contributors to connectivity-based predictive models or “fingerprints” of cognitive processes [Rosenberg et al., 2016] or psychopathology [Elliott et al., 2017]). Finally, a recent attempt to quantify the utility of parcellations of varying granulatiy for connectivity-based predictive modelling have shown fine grained parcellations lead to higher predictive accuracy [Li and Gowtham, 2018].

Important questions that can be examined at a finer level of analysis are 1) Do within- and between-network connections show similar levels of genetic and environmental influences? 2. Are regions of a particular resting-state network homogeneous in terms of their pattern of genetic and environmental influences across all connections? 3. What are useful applications of high resolution genetic mapping? For example, can differences between regions’ patterns of genetic connectivity be used to generate hypotheses about or explain differences in their functions? Answering these questions speaks to recent efforts to “carve nature at its joints” and thus has important implications for how we conceptualize resting state connectivity as a biomarker or candidate endophenotype for behaviors of interest.

To answer these questions, we analyzed resting state data from two comparably-aged adult twin samples: the Colorado Longitudinal Twin Study (LTS; N = 465, including 102 complete monozygotic [MZ], 91 complete same-sex dizygotic [DZ] pairs, and 79 singletons), and the Human Connectome Project (HCP; N = 422, including 136 complete MZ and 75 complete same-sex and opposite-sex DZ pairs). The inclusion of both samples allowed us to examine replicability of general patterns rather than significance of specific effects. We decomposed the functional connectome of each individual into pairwise correlations between 264 individual regions (referred to as connections) from a widely used and independently derived brain parcellation ([Power et al., 2011]; i.e., 34,716 functional connections). In comparison to coarser parcellations, this parcellation was developed to reflect functional distinctions between small parts of cortex [Wig et al., 2011], is accompanied by metadata assigning each region to one of 14 function-specific resting-state communities (e.g., visual network, default network, etc.), and is within a window of optimum dimensionality that maximizes reproducibility [Thirion et al., 2014].

We addressed the primary questions-of-interest by applying a classic univariate twin model to each connection (see **Materials and Methods - Genetic Models**) to estimate the proportion of variance in connection strength explained by additive genetic influence (A or heritability; the sum of a large number of genetic variants that additively influence a trait), shared environmental influence (C; influences that increase similarity of siblings), and non-shared environmental influence (E; influences that decrease similarity of siblings). The resulting high-resolution genetic and environmental maps allowed us to investigate differences between within-network and between-network connections (*question 1*) and also investigate the distribution and patterns of genetic influence for regions of a given *a priori* resting state network (*question 2*).

As a prelude to the results, we find substantial heterogeneity of genetic influence in the connectome. Our third question-of-interest broadly asks how these results can be utilized in the areas of cognitive and clinical neuroscience, so we provide three example applications. First, we show that although analysis of the default network as a whole (as has been done in the past) is equivalent to averaging the analysis of many default network connections (as done in the current study), there is important detail lost in the averaging. That is, certain hyper-localized parts of the default network have very strong genetic influence while others have approximately zero genetic influence. This method could be applied, in the future, to different pieces of the connectome to discover novel classifications of brain areas within less studied networks or to generate hypothesis about discriminant functions of brain areas. Second, we show that genetic influence on local connections are separable from genetic influence on networks as a whole by utilizing bivariate genetic models capable of quantifying common and unique genetic influences on variation in multiple measures. Finally, we simulate many connectivity-based predictive models and estimate the heritability of these signals using both low-resolution (network) and high-resolution (region-based) heritability estimates. This analysis reveals systematic biases that arise when using heritability estimates from network-based connectivity measures to explain the etiology of high-resolution predictive model weights. Together, these analyses elucidate differences in the genetic and environmental etiology of connections of different type/function and demonstrate possible applications in a variety of domains.

## Results

### Group Average Connectomes

Visual comparison of mean phenotypic connectivity matrices for each sample to one another and to matrices reported in prior work using independent samples (e.g., Figure 3 of [Cole et al., 2014]; Figure 2 of [Reineberg and Banich, 2016]) reveals striking similarity, especially in the prominence of resting-state networks along the diagonals (**Figure S1** [LTS] and **Figure S2** [HCP]).

### Univariate Twin Models

Connection-wise estimates of additive genetic influence for the LTS and HCP samples are shown in **Figure 1a, b** (lower triangles). In the LTS sample, additive genetic influence was moderate and bimodally distributed across the connectome such that 16,317 of 34,716 unique connections were estimated as having zero heritability while a separate, positively-skewed distribution described the heritability of 18,399 connections (M = 0.112, SD = 0.078, Skew = 0.795, Min = 0.001, Max = 0.456). Similarly, for the HCP sample, 13,516 of 34,716 unique connections were estimated as having zero heritability while a separate, positively-skewed distribution described the heritability of 21,200 connections (M = 0.127, SD = 0.082, Skew = 0.608, Min = 0.001, Max = 0.504).

Shared environmental influences generally explained less variance than genetic influences, as shown in **Figure 1a, b** (upper triangles). In the LTS sample, shared environmental influence was weak to moderate and bimodally distributed across the connectome such that 20,806 of 34,716 unique connections were estimated as having zero shared environmental influence while a separate, positively-skewed distribution described the shared environmental influence of 13,910 connections (M = 0.081, SD = 0.057, Skew = 0.811, Min = 0.001, Max = 0.400). Similarly, for the HCP sample, 20,297 of 34,716 unique connections were estimated as having zero shared environmental influence while a separate, positively-skewed distribution described the shared environmental influence of 14,419 connections (M = 0.093, SD = 0.064, Skew = 0.720, Min = 0.001, Max = 0.367). Although C estimates were low, there is evidence of moderate shared environmental influence in several pieces of the connectome such as within-default network connections, within-sensory somatomotor connections, and default-to-other connections.

In both samples, non-shared environmental influences were high across the entire connectome (MLTS = 0.908, SDlts = 0.081; Mhcp = 0.884, SDhcp = 0.086) and negatively skewed (Skew_LTs_ = −0.770, Skew_HCP_ = −0.513). Connection-wise estimates of non-shared environmental influences are shown in the lower and upper triangles of **Figure S3** for the HCP and LTS samples, respectively. It should be noted that E estimates include measurement error, so reliability of connections was tested for the HCP sample and found to be high (M = 0.849, SD = 0.061; see **Supporting Information - Reliability**), suggesting that the high E estimates across the connectome are unlikely to solely reflect random measurement error.

In general, both sample had very similar patterns of heritability. To be sure any sample differences in estimates was due to true differences between the two samples, we compared 6 to 30 minutes of data within the HCP sample. This analysis leads us to believe the differences between the LTS and HCP samples are related to sample differences rather than data quantity. When comparing 6 minutes to 30 minutes of HCP data, heritability estimates changed in magnitude (mean h^2^_6 min_ = 0.088, mean h^2^_30 min_ = 0.127) but remained similar in pattern (i.e., were correlated with one another). See **Supporting Information - 6 vs. 30 Minutes of Resting Data** for more informations about this analysis.

**Figure 1.**
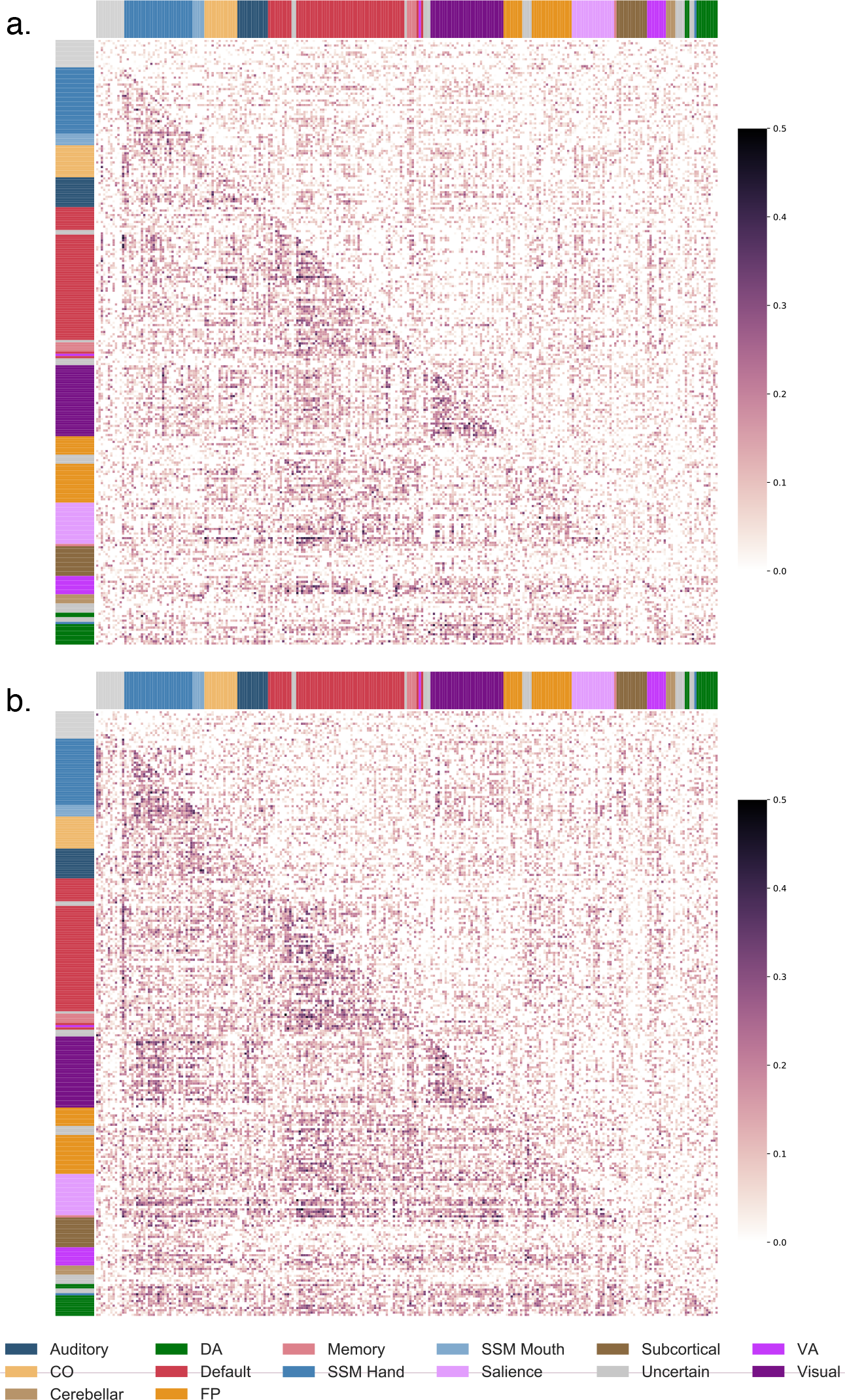
Connection-wise estimates of additive genetic (A) and shared environmental (C) influences. Matrices contain estimates from univariate twin models with the spatial location of each cell (estimate) corresponding to the functional connection between two regions. Assignment to *a priori* networks is represented by colored bars along x and y axes. Because triangles are redundant, different estimates are displayed in upper and lower triangles. a. LTS sample A and C estimates are in the lower and upper triangles, respectively. b. HCP sample A and C estimates are in the lower and upper triangles, respectively. CO = cingulo-opercular, DA = dorsal attention, FP = frontoparietal, SSM = sensory/somatomotor, VA = ventral attention.

### Within- and Between-Network Connections

To examine whether high resolution mapping of genetic influence reveals differences in within- versus between-network connections, (*question 1*), we investigated heritability estimates for connections of those types. First, we considered average heritability across all connections considered to be within the same *a priori* network versus all between-network connections. In both samples, within-network connectivity was more heritable than between network connectivity (**Table S1a**; whole connectome results). This effect was present even when controlling for the estimated test-retest reliability of each connection in the HCP sample (see **Supporting Information - Reliability**).

We also quantified differences in heritability for within and between network connections for each *a priori* network individually, as shown in **Figure S4**. In both samples, within-network connections tended to be more heritable on average than between-network connections. In both samples, the default, sensory-somatomotor hand, auditory, dorsal attention, visual network had significantly higher heritability for within- than between-network connections (**Table S1b** and **Figure S4**), but the subcortical and uncategorized networks had significantly higher heritability for between- than within-network connections. Although within-network connections tended to be more heritable on average than between-network connections, the distributions of between-network connections tended to be more positively skewed, perhaps suggesting there are a minimal number of highly heritable between-network connections.

Finally, we quantified differences in heritability for within- and between-network connections at the level of regions (each of the 264 regions of the parcellation). Of regions that had significantly higher heritability for within- than between-network connections in the LTS (n = 90) and HCP (n = 86) samples, 45 regions showed the effect in both samples (See **Table S2**). These regions were from the sensory-somatomotor (7), default (19), and visual (15) networks, as well as one regions from each of the frontoparietal and auditory networks. One region from salience network and one unclassified region had significantly higher heritability for between-network connections than within-network connections.

### Clustering Reveals Large Genetic Communities

Given the heterogeneity in genetic influence described above, we wanted to explore whether the best way to describe genetic communities in the connectome was in terms of *a priori* functional networks. The distribution of A estimates within *a priori* networks (**Figure S4**) suggests regions of any given resting-state community have a variety of different patterns of heritable connectivity across the connectome. Variation in the patterns of genetic influence for the different regions of each network can be explored with clustering analysis, which seeks to group regions together based on how similar their pattern of heritable connectivity is. Thus, we conducted a data-driven clustering analysis to group together regions with similar patterns of heritable connectivity. K-means clustering groups rows of the additive genetic influence matrix (**Figure 1**) that show similar patterns of heritable connectivity with all other regions. This analysis could reveal that the 264 regions cluster together in a manner similar to *a priori* networks or in a novel way (e.g., a cluster of regions with highly heritable connectivity to some default and frontoparietal network regions, but minimally heritable connectivity to other regions). We analyzed average silhouette scores for clustering solutions (i.e., *k*-values) from 2 to 20, shown in **Figure 2a**, and discovered stable solutions at *k*-values of 2, 3, 4, 7, and 15 in the LTS sample and *k*-values of 2, 3, 7, and 13 in the HCP sample.

Because the first stable solution at a *k*-value of 2 primarily differentiated regions based on magnitude of heritability (i.e., more heritable vs. less heritable regions), we explored the *k* = 3 clustering solution first. This level provides the highest level overview of patterns of genetic influence across the connectome. The three clusters from the *k* = 3 stable solution will be referred to as super-clusters throughout the remainder of the manuscript. The 3-cluster solution for the LTS sample is shown in **Figure 2b**. **Figure 3** provides an overview of both the spatial location of the regions in each super-cluster (a-c) and also the composition of those superclusters in terms of the regions assignments to *a priori* networks (right-most column).

Overall, the results revealed that regions cluster at a level superordinate to *a priori* notions of resting-state community structure. Super-cluster 1 was composed of 135 regions from all *a priori* networks with no distinct pattern of heritable connectivity. Generally, heritability was low to moderate for all connections in these 135 regions. Super-cluster 2 regions had especially heritable connectivity to pure sensory and sensory-somatomotor regions as well as moderately heritable connectivity to regions from cognitive networks (default, cingulo-opercular, frontoparietal, and attention networks). Super-cluster 2 was composed of 39 regions from a variety of pure sensory (e.g., visual) and sensory-somatomotor networks. Super-cluster 3 regions had especially heritable connectivity to default, frontoparietal, salience, dorsal attention, and visual network regions. Super-cluster 3 was composed of 90 regions that can best be summarized as the majority of the default network as well many fronto-parietal regions, among others. The *k* = 3 solution of the LTS sample maps closely on to the *k* = 3 solution of the HCP sample with only minor difference (discussed further in **Supporting Information - Clustering**).

Higher-order clustering solutions from the HCP sample give insight into how these large genetic communities break down into more specific patterns of genetic influence. **Figure S5** shows the composition of sub-clusters from the 15-cluster solution. Some of the sub-clusters of the 15-cluster solution remain large and highly heterogeneous (i.e., composed of regions from many different *a priori* resting-state communities). Others are smaller and relatively pure, such as sub-clusters 5, 6, 8, 9, 10, 11, 13, and 15, which contain one to three types of regions. Of note is that regions from most *a priori* resting-state communities split into several different clusters, supporting the conclusion that *a priori* networks contains several sets of regions that have unique patterns of heritable connectivity across the connectome. For example, default network regions can be found in 10 of the 15 sub-clusters. Future work could explore this and other higher-dimensionality clustering solutions (as we limited our clustering analysis to between 2 and 20 clusters) to possibly reveal novel communities of brain regions.

**Figure 2.**
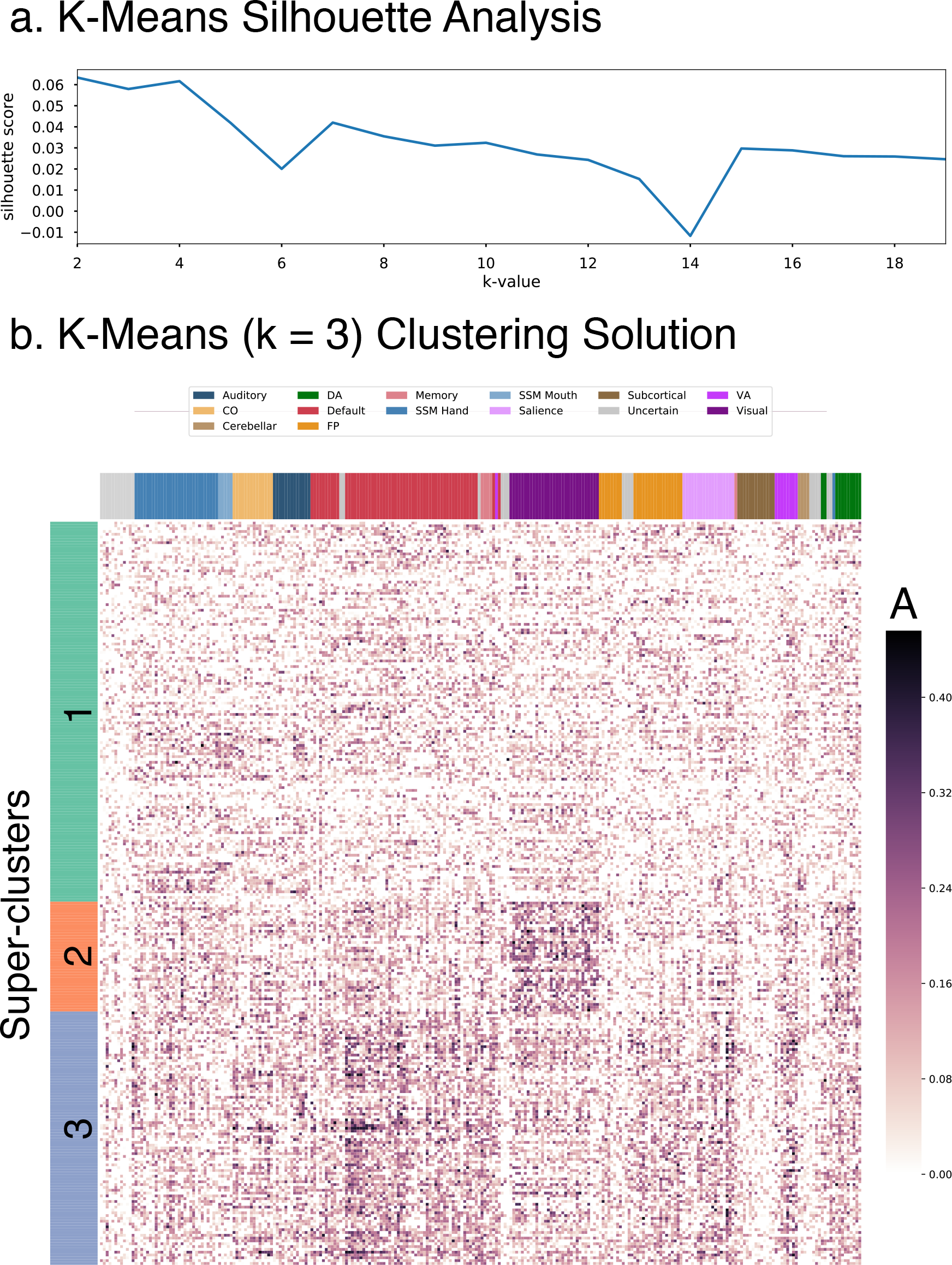
K-Means 3-cluster solution. Row-wise clustering of LTS additive genetic (A) estimates reveals several stable clustering solutions of retions with similar patterns of connectivity heritability. Super-clusters (*k* = 3) are described in detail. a. Silhouette analysis revels stable clustering solutions at *k*-values of 2, 3,4,7, and 15. b. Clustered version of LTS A estimates for *k* = 3 solution. CO = cingulo-opercular, DA = dorsal attention, FP = frontoparietal, SSM = sensory/somatomotor, VA = ventral attention.

## Applications

### Revealing particularly heritable connections bridging two resting-state networks

We performed a series of follow up analyses in the LTS sample on an example set of between-network connections to explore heterogeneity of genetic influence in a more focused system. Prior work has found the most heritable default network connection is the connection to the sensory/somatomotor network [Yang et al., 2016]. We extracted the many connections between default network regions and sensory/somatomotor regions for a more detailed investigation (n = 2,030, all pairwise connections between 35 sensory/somatomotor and 58 default network regions). **Figure 4a** shows distribution of nonzero heritability estimates for default-to-sensory/somatomotor connections (Max h^2^ = 0.360). Additionally, we estimated heritability for the default-to-sensory/somatomotor network connection as a whole using network templates from a popular network parcellation (Yeo et al, 2011; h^2^ = 0.240). The network-derived heritability estimate is shown as a dotted vertical line in **Figure 4a**.

The fine-grained analysis reveals a subset of default to sensory/somatomotor connections have moderate genetic influence, whereas many have minimal to no genetic influence. **Figure 4b** shows the most heritable of the default to sensory/somatomotor connections. We find the most heritable connections are between superior, medial frontal cortex and the sensory/motor strips; hub regions of the default network and the sensory/motor strips; as well as connections between the middle temporal lobes and the sensory/motor strips. A recent meta-analysis of thousands of functional MRI studies revealed the function of many of these superior, medial regions is related to “conflict”, “working memory”, and “inhibition” [de le Vega et al., 2016]. The hubs of the default network have an important role in the valuation of motivationally-salient and personally-significant information [Andrews-Hanna, 2011]. The middle temporal gyrus is part of a default network subsystem responsible for introspection about mental states (Ibid.). Here we have identified particularly strong heritability for connectivity between these places and sensory/somatomotor regions. Future work could explore other network connections of interest in the same way as we have here to reveal other novel characterizations of between-network connections. In summary, studies that utilize network-derived estimates should note they may be averaging many connections with heterogeneous genetic influence, which can result in a network-derived heritability estimate that is an underestimate of the maximum heritability between smaller functional units. Important distinctions between connections with particularly strong heritability would be missed in network-based studies.

### ROI-based Connectivity Estimates are Genetically Separable from Network-derived Estimates

Variation in heritability estimates does not imply separate genetic influences (i.e., sets of genes responsible for the difference in heritability). For example, the network estimates quantified in earlier work could be driven by the same or different genetic variants as the connections in the current report. Here we performed a bivariate genetic analysis (models pictured in **Figure 6** and described in **Methods - Genetic Models**) between the many default-to-sensory/somatomotor connections described above and the network-derived connectivity estimate for these two networks as a whole. Whereas the univariate models described above quantify genetic influences on local connections alone, these bivariate analyses quantify the degree to which local functional connectivity is genetically separable from the network-derived estimate of connectivity between the default and sensory/somatomotor networks.

First, we found local and network connectivity does have strong genetic correlation (“rA”; pictured in **Figure 5a**; M_|*rA*|_ = 0.790, SD_|*rA*|_ = 0.324, Min = −1.000, Max = 1.000) as would be expected given the network-derived connectivity estimate includes the same time series data as the more focused 2,030 connections. However, we find there is residual genetic influence not accounted for by the network-derived estimate (pictured in **Figure 5b**; M = 0.051, SD = 0.076, Max = 0.267). Together, these analyses demonstrate the local connections have unique genetic influences although there is certainly a substantial, common genetic component captured when utilizing network-derived connectivity estimates.

Genetic correlations between individual connections and higher-level measures of individual differences in connectivity may be of interest to those invested in graph theoretic analysis of the brain. Graph theory analysis offers many possible summary measures of connectivity and seeks to summarize the brain in the context of complex network dynamics - for example, the degree to information can be shared amongst distributed brain systems (i.e., integration as measured by global efficiency; for review, see [Rubinov and Sporns, 2010]). In **Supporting Information - Bivariate Analysis** we describe an analysis in which we quantify the degree to which connections are genetically separable from global efficiency. We find connections have residual genetic influence not accounted for by genetic influences on global efficiency.

**Figure 3.**
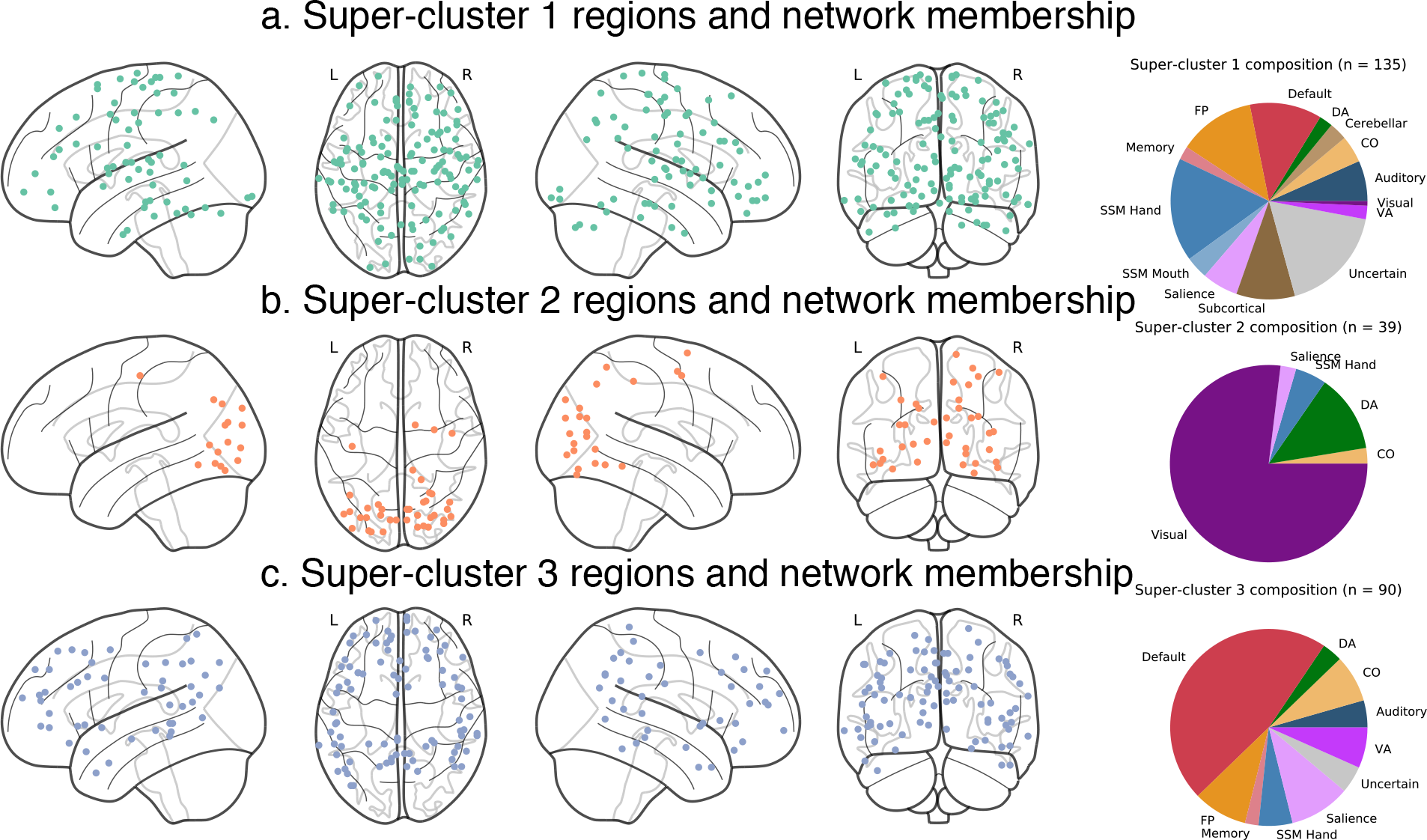
K-Means 3-cluster summary. Spatial location of regions from LTS super-clusters 1-3 of *k* = 3 solution. a. Super-cluster 1 regions were widely distributed across the brain. b. Super-cluster 3 regions were located primarily in sensory and somatomotor areas. c. Super-cluster 2 regions were located across lateral prefrontal, lateral parietal, mid and anterior temporal, midline frontal, and cingulate areas. n = number of regions, CO = cingulo-opercular, DA = dorsal attention, FP = frontoparietal, SSM = sensory/somatomotor, VA = ventral attention.

### Investigating Genetic Correlation of In-scanner Movement and Connectivity

Prior work has found in-scanner movement, as measured by mean frame displacement, is heritable (h^2^ = 0.313 - 0.427, [Hodgson et al., 2016]). In all analyses reported above, we controlled for movement via single subject denoising and summary movement covariates, so the results do not reflect covariance with movement (i.e., are equivalent to a “specific heritability” estimate as described in the bivariate analyses presented previously). However, we wondered whether or not some connections (before regressing out summary movement covariates) might be genetically related to movement. Specifically, in the LTS sample, we utilized bivariate genetic models (**Figure 6** and **Methods - Genetic Models**) to quantify where in the connectome there is overlapping genetic influence between connectivity (without summary movement covariates) and in-scanner movement (i.e., genetic correlations or “rA”). In an initial univariate analysis, we found translation (average motion in the x, y, and z planes) was not heritable (h^2^ = 0.012) and weakly influenced by shared environmental influences ((C^2^ = 0.168). Rotation (average roll, pitch, and yaw movements) was highly heritable (h^2^ = 0.884). When combined into a single composite summary (i.e., average of translation and rotation), motion was moderately-to-highly heritable (h^2^ = 0.578).

Only rotation movement had a genetic component, so we subjected rotation movement to the bivariate analysis pipeline. Genetic correlations between rotation movement and connections were moderate-to-strong (M_|*rA*|_ = 0.473, SD_|*rA*|_ = 0.381, Min = −1.000, Max = 1.000). The matrix of genetic correlations is presented in **Figure S9**. Interestingly, there are portions of this matrix with strong positive genetic correlations indicating there may be common genes that are associated with level of movement and connectivity strengths (or reflecting that movement may induce artifactual connectivity differences in these places). These include some cerebellar connections (such as those to the default network) and connections of regions with no functional network assignment. Within-network connections seem to have the weakest genetic correlations between movement and connectivity strength, however, within-sensory/somatomotor network connections may be an exception to that observation. Negative genetic correlations were less common, but visual network connections may be enriched for negative genetic correlations, especially connections to the auditory and salience networks. Future work should investigate these genetic correlations in more detail, perhaps mapping movement-connectivity genetic correlations onto existing models of movement-related susceptibility for connections of different type and distance. In summary, had we not residualized connectivity with regard to movement in the main analyses of the current study, we likely would have interpreted large heritability estimates across the entire connectome as meaningful when, in fact, they are driven partially-to-mostly by strong genetic influence on rotation movement.

**Figure 4.**
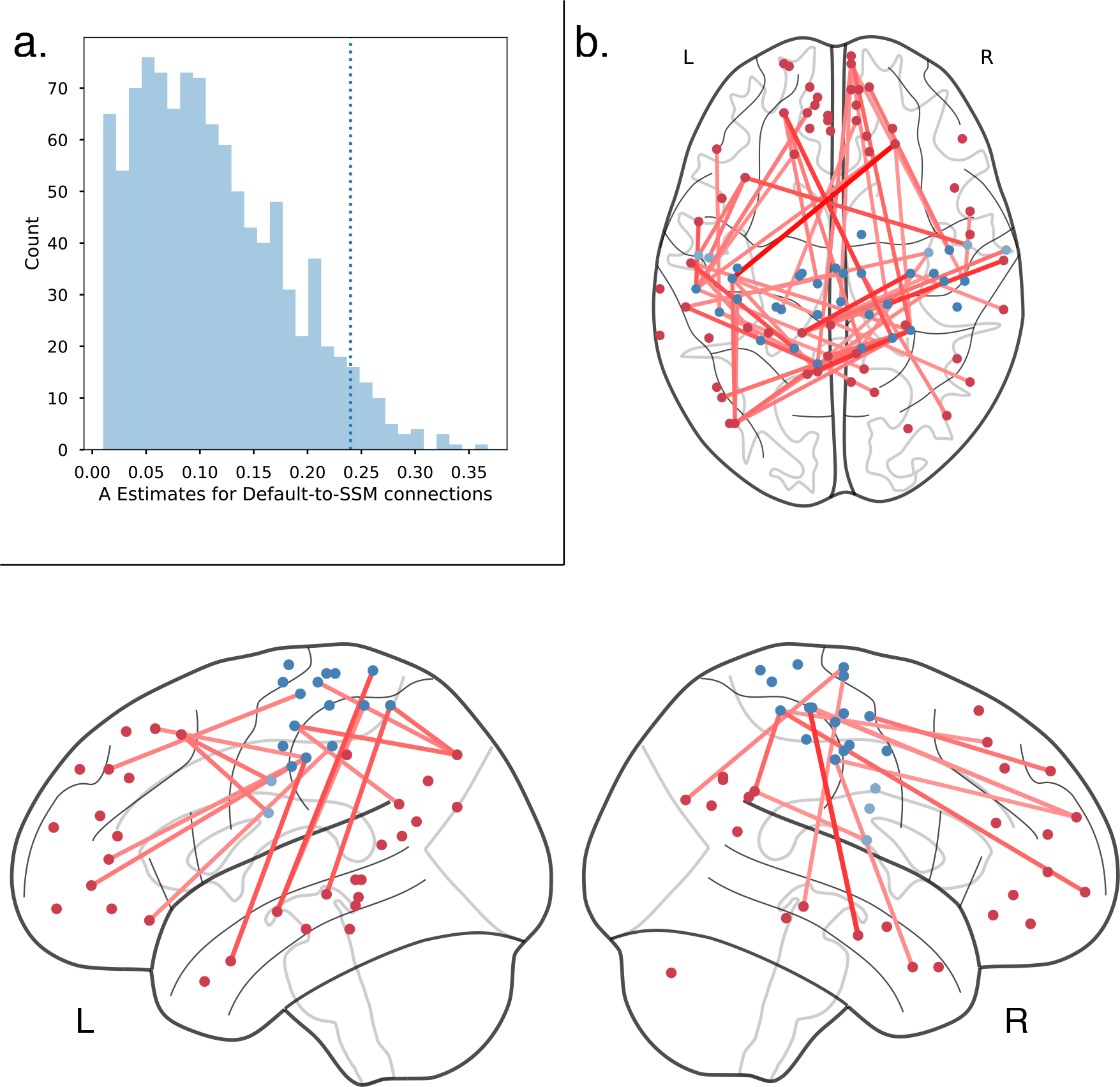
Default-to-Sensory/somatomotor Connections and Heritability. a. Distribution of nonzero heritability estimates for default-to-sensory/somatomotor connections. Dotted line (h^2^ = 0.240) indicates the heritability estimate for the connection between the default and sensory/somatomotor network when analyzed as a whole rather than many connections between smaller, focal regions-of-interest. b. Connections between default (red) and sensory/somatomotor (blue) regions with heritability higher than network-derived heritability estimate (h^2^ > 0.240) projected onto the brain. The most heritable connections between the default and sensory/somatomotor network are between the sensory/motor strips and superior, medial frontal, posterior cingulate, and middle temporal regions of the default network.

**Figure 5.**
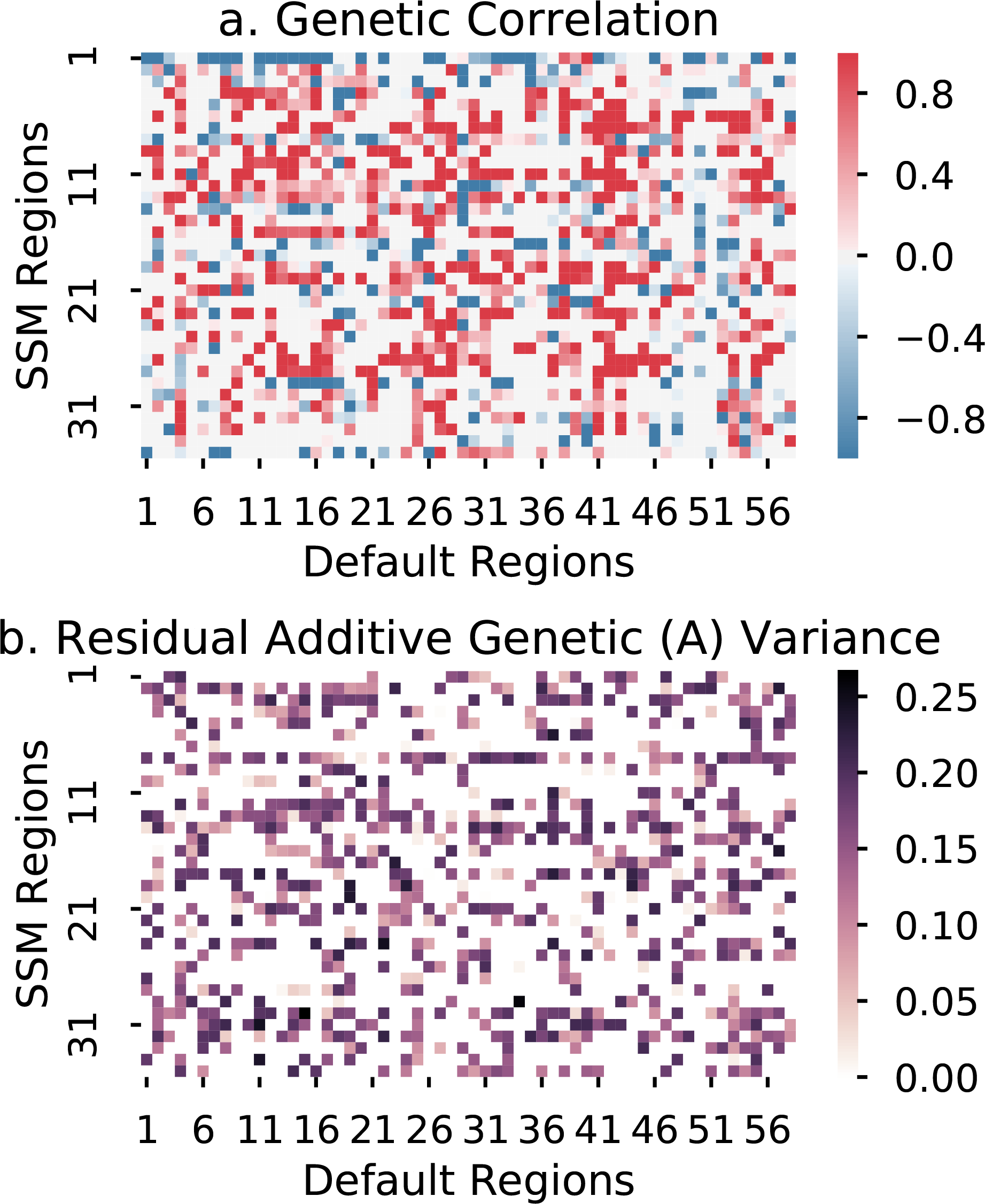
Default-to-Sensory/somatomotor Connections - Bivariate Analysis. a. Genetic correlation between network-derived estimate of default to sensory/somatomotor connectivity and many default to sensory/somatomotor connections b. Residual additive genetic influence on many default to sensory/somatomotor connections after accounting for genetic variance shared with a network-derived estimate of default to sensory/somatomotor connectivity.

## Discussion

Across all analyses, we found converging evidence of etiological heterogeneity in the functional connectome. High-resolution mapping reveals a distribution of genetic and environmental influence that may be missed by approaches that summarize functional connectivity at the level of larger ROIs, networks, and global summary measures of the connectome. More specifically, we found differences in genetic influences for connections of different type (i.e., higher heritability of connections between regions of the same functional network versus between regions of different functional networks). This pattern was present across the whole connectome and especially for the default, sensory/somatomotor, dorsal attention, and visual networks. This result provides preliminary evidence that the organization of the brain into networks based on function may be driven by genetic influences on connections between regions involved in the same processes. Interestingly, prior work has established specific patterns of gene expression within functional networks [Richiardi and Altmann, 2015], a possible mechanism linking these observations of genetic influence to specific functions. Importantly, we showed how high resolution heritability estimates might be used to define novel communities of regions based on there pattern of genetic influences as well as how to isolate pieces of the connectome with particularly high genetic influence (i.e., may be candidate endophenotype). Finally, we showed genetic influences on connections are separable from genetic influences on network connectivity and in-scanner movement during the resting-state scan (a frequently-discussed source of nuisance signals in functional imaging studies).

Although a number of our analyses were summarized by *a priori* functional networks, the broad range of genetic estimates across the connectome led us to question whether alternative groupings could better describe patterns of heritability in the connectome. A clustering procedure revealed a novel finding: Regions grouped together based on patterns of heritable connectivity at a level that was superordinate to that of classic resting-state communities. In both samples, we found stable super-clusters of regions. Most notably, a set of “cognitive” and “sensory/somatomotor” regions had characteristic patterns of highly heritable connections to regions of higher-level (e.g., default network) and lower-level (e.g., visual) functions, for the cognitive and sensory/somatomotor clusters respectively. Although the description of the these super-clusters as higher and lower-level is likely an oversimplification, it is a worthwhile descriptive tool until future work dissects the role of these sets of brain regions. In both samples, we found evidence of a super-cluster of regions with very consistent low-to-moderate heritability to all regions (i.e., no distinct features) and high non-shared environmental influence. Analyses of connection-wise reliability (very high in all connections in the HCP sample) suggest that these non-shared environmental influence estimates do not simply reflect random measurement error. Thus, future work should seek a more thorough understanding of the environmental factors influencing this nondescript set of regions. Stable clustering solutions were also found at levels of granularity similar to classic resting-state communities, but, interestingly, these genetic clusters were quite dissimilar to the *a priori* networks. Notably, in our example 15 cluster analysis, regions from the default network broke into several sub-clusters which were differentiated on heritability of connectivity to other default network regions and to regions of other networks such as the frontoparietal network. Future work should dissect these finer-grained parcels in more detail, especially the degree to which the represent novel or previously-described communities. Finer-grained, stable clustering solutions could be explored in more detail too as those may reveal small communities with highly characteristic patterns of heritable connectivity which may not align with known clusters of regions based on community detection performed on phenotypic functional connectivity.

Our use of bivariate genetic models represents a substantial development in neuroimaging genetics. We found local connections showed genetic influences independent from genetic influences on network connectivity, in-scanner movement, and a global summary measure of the brain. Residual genetic influence justifies analysis at the level of small regions and is an important commentary on an ongoing debate about the proper level of analysis of connectivity, suggesting all levels may be complementary. A practical application of this evidence of residual genetic influence would be to the study of multivariate functional connectivity signals as a predictor of individual differences in some cognitive ability or clinical variation (i.e., connectivity-based predictive modelling,“fingerprinting”, or “connecto-typing”). Our results suggest influence on a whole connectome signal will be diverse and not accurately represented in network-based or global summary measures of the connectome. Specifically, our results show network-derived estimates could over- or underestimate heritability of different pieces of a predictive model by assigning the network-derived measure to all connections of the same type, which we have shown here to have a distribution of genetic influence and are genetically separable from the network-derived measure of connectivity. Regarding movement, we showed evidence of both overlapping and distinct genetic influences for in-scanner movement and connections in an analysis which utilized connectivity estimates that did not control for individual variation in movement. This analysis supports the strong emphasis in the resting-state literature to adequately control for nuisance signals associated with in-scanner movement. Additionally, these results are the first, to our knowledge, make a clear distinction between the etiology of various types of in-scanner movement. We found translation movement was not heritable but rotation movement was highly heritable.

Although we opted to utilize bivariate genetic models for the purpose of clarifying the specificity of genetic effects with regard to nuisance signal and broader measures of connectivity, future work could apply the same bivariate analysis to the relationship between local connections and clinical symptomatology and/or cognitive abilities. Such an application could identify novel brain-based candidate endophenotypes or focus intervention studies to novel locations. A similar approach has been used in the neuroanatomical/clinical endophenotyping literature in which bivariate genetic models have been used to identify the genetically-influenced neurobiological underpinning of disorders such as major depressive disorder [Glahn et al., 2012] and the genetically-influenced neurobiological underpinnings of schizophrenia that are shared with other psychiatric disorders ([Lee et al., 2016]).

Our approach is not without caveats. Although we highlighted overlapping results in both samples, we also observed some sample-specific results. Notably, shared environmental influence was lower across the entire connectome in the LTS sample than in the HCP sample. Although it is not unusual to find a lack of shared environmental influence (e.g., in the anatomical MRI [Eyler et al., 2011] and cognitive literatures), sample differences could be due to reliability differences in the measurement of resting-state functional connectivity (e.g., scan time of 30 minutes [HCP] versus 6 minutes [LTS] is known to produce more reliable results [Gordon et al., 2017] although we did find 6 minutes of HCP resting data produced heritability estimates very similar to those produced using 30 minutes of data) and/or due to demographic differences in the two samples (e.g, the Colorado sample is less racially diverse and sampled from higher socioeconomic status communities than the HCP sample). Socioeconomic status differences could certainly explain sample differences in the current study given prior work showing elevated shared environmental influence on variation in IQ for individuals near or below the poverty line [Turkheimer et al., 2003]. Effects of SES in a subset of the HCP sample have been partially explored previously and shown to influence brains connectivity [Smith et al., 2015]. Regarding modeling of genetic influences, a small literature suggests classic twin modeling procedures may bias estimates (upward in the case of A and downward in the case of E) when compared to models that do not impose boundary constraints on parameters [Carey, 2005]. Future work should compare these approaches and report notable differences, if any, in the genetic profile of affected connections.

Overall, we demonstrate the utility of fine-grained A, C, and E estimates by showing the genetic organization of the brain is diverse and not as one would expect based solely off the classic functional organization of the phenotypic connectome. Our analysis sits in a continuum of dimensionality reductions that spans multiple levels of brain organization (i.e., from global summary measures to voxels), so, obviously one must ask if genetic neuroimaging studies should continue to assess the etiology of finer grained parcellations in the future. Our demonstration of residual genetic variance for local connections in the bivariate analyses certainly demonstrates the added value of a fine-grained approach in addition to a single summary measure of the connectome. But, our results also suggest a trade-off between reliability and interpretability/application: Large networks maximize heritability estimates but are of imprecise function and cannot be used to dissect the etiology of highly dimensional signals that are most useful for predictive modeling. Parcellations in the range of 200-500 might be recommended for region-based approaches in the future because there are numerous well-vetted atlases [Power et al., 2011, Craddock et al., 2012, Gordon et al., 2016] designed to differentiate homogeneous functional brain units while maximizing reliability (which could become an issue in voxel-based approaches). There is still room for determining the best functional parcellation scheme among these possible alternatives, with genetic etiology as one possible mechanism for evaluating the quality/usefulness of the parcellations. In conclusion, our approach has important implications for investigations of neuroimaging-based biomarkers by 1. quantifying which pieces of the connectome are heritable and thus can be investigated as a potential endophenotype or marker of genetic risk; 2. serving as a model for future studies seeking a greater understanding of a broad literature of traits; and 3. establishing the foundation of a taxonomy of functional connections based on genetic influence.

## Materials and Methods

The current study is a parallel analysis of resting-state data from a sample of adults recruited from the Colorado Longitudinal Twin Study (referred to as LTS throughout the manuscript) and 422 adults from a publicly available data set from the Human Connectome Project (referred to as HCP throughout the manuscript).

### Participants

Participants from the LTS sample were 465 individuals (Age Range = 28 - 32, M_age_ = 28.7 years, SD_age_ = 0.63 years; 199 males) after 35 participants were removed due to excessive movement during the scanning session based on the criteria of greater than 2 mm translation (motion in X, Y, or Z plane) or 2 degrees rotation (roll, pitch, or yaw motion) (*n* = 34), and failure of the presentation computer to display a fixation cross during the resting scan (*n* = 1). Of the 465 individuals, there were 102 pairs of MZ twins, 91 pairs of DZ twins, 34 MZ twin singletons, and 45 DZ twin singletons. Singletons are members of twin pairs whose co-twins either did not participate or were excluded from analysis.

Singletons were only used in the genetic analyses to calculate connectivity means and variances. All participants were recruited from the Colorado Longitudinal Twin Study which recruited from the Colorado Twin Registry based on birth records. Comparisons with normative data on several measures suggests that the sample is cognitively, academically, and demographically representative of the state of Colorado. Based on self report, the LTS sample is 92.6% White, 5.0% âĂIJmore than one raceâĂİ, <1% American Indian/Alaskan Native, <1% Pacific Islander, or did not report their race (1.2%). Hispanic individuals composed 9.1% of the sample. Additional information about the LTS sample can be found elsewhere [Rhea et al., 2006, Rhea et al., 2013]. Participants were paid $150 for participation in the study or $25 per half hour for those who did not finish the entire three-hour session. The study session involved the administration of behavioral tasks that measured EF ability as well as acquisition of anatomical and functional brain data via magnetic resonance imaging.

HCP participants were 422 individuals (M_age_ = 29.2 years, SD_age_ = 3.46 years, Range = 22 - 35 years; 171 males) selected from the most recent HCP data release because they were part of complete pairs of twins who completed the anatomical and functional imaging components of the study. This subset of HCP participants were 136 MZ pairs and 75 DZ pairs with race reported as 82.7% White, 11.3% Black/African American, 4.5% Asian/Nat. Hawaiian/Other Pacific Is., and <1% Unknown/Not reported, More than one, or Am. Indian/Alaskan Nat. each.

### Procedure

Testing took place in a single three-hour session. Following review and obtainment of informed consent, participants were familiarized with the imaging procedures. If both twins of a pair participated on the same day, the twins completed the protocol sequentially (twin order randomized) with the same ordering of behavioral testing and imaging acquisition. The resting-state scan always occurred first in the imaging protocol, before tasks. All study procedures were fully approved by the Institutional Review Board of the University of Colorado Boulder. All participants read and agreed to the informed consent document prior to their initial enrollment in the study and at each follow-up assessment. Testing for the HCP sample participants has been explained thoroughly in prior work [Van Essen et al., 2013].

### Brain Imaging

Participants from the LTS sample were scanned in a Siemens Tim Trio 3T (n = 250) or Prisma 3T (n = 215) scanner. Neuroanatomical data were acquired with T1-weighted MP-RAGE sequence (acquisition parameters: repetition time (TR) = 2400 ms, echo time (TE) = 2.07, matrix size = 320 × 320 × 224, voxel size = 0.80 mm × 0.80 mm × 0.80 mm, flip angle (FA) = 8.00 deg., slice thickness = 0.80 mm). Resting state data was acquired with a 6.25 minute T2*-weighted echo-planar functional scan (acquisition parameters: number of volumes = 816, TR = 460 ms, TE = 27.2 ms, matrix size = 82 × 82 × 56, voxel size = 3.02 mm × 3.02 mm × 3.00 mm, FA = 44.0 deg., slice thickness = 3.00 mm, field of view (FOV) = 248 mm). During the resting-state scan, participants were instructed to relax and stare at a fixation cross while blinking as they normally would. Scanner was included as a nuisance regressor in all analyses involving the LTS dataset. Resting-state acquisition in the HCP sample is described in detail elsewhere [Smith et al., 2013], but briefly, each participant completed an anatomical and four, 15-minute resting state scans (eyes fixated) in the context of a large imaging and behavioral testing battery. In the current study, the first two 15-minute resting state scans were utilized.

### Preprocessing and Connectome Extraction

All processing of LTS brain data was performed in a standard install of FSL build 509 [Jenkinson et al., 2012]. To account for signal stabilization, the first 10 volumes of each individual functional scan were removed, yielding 806 volumes per subject for additional analysis. The functional scans were corrected for head motion using MCFLIRT, FSL’s motion correction tool. Brain extraction (BET) was used to remove signal associated with non-brain material (e.g., skull, sinuses, etc.). FSL’s FLIRT utility was used to perform a boundary-based registration of each participant’s functional scan to his or her anatomical volume and a 6 degree of freedom affine registration to MNI152 standard space. To account for motion and other noise signals known to pollute resting-state analyses, LTS scans were subjected to AROMA, an automated independent components analysis-based, single-subject denoising procedure [Pruim et al., 2014]. Signal was extracted from masks of the lateral ventricles, white matter, and whole brain volume and regressed out along with a set of 6 motion regressors and associated first and second derivatives. Finally, the scans were band-pass filtered (.001 - .08 Hz band).

Preprocessing for HCP data is described elsewhere [Glasser et al., 2013]. Briefly, HCP scans were subjected to minimal preprocessing and FIX, a semi-automated single-subject denoising procedure [Salimi-Khorshidi et al., 2014]. Additionally, we regressed out the mean greyordinate time series from each scan as a proxy for the global signal (as suggested by [Burgess et al., 2016]). HCP scans were bandpass filtered (0.001 - 0.080 Hz band).

For each participant, we extracted the BOLD time series from each of 264, 1 cm spherical ROIs, drawn from [Power et al., 2011], which serve as the nodes for the present analysis. This analysis was performed in volume space to maximize similarity between the two samples. We used these nodes as they are drawn from a meta-analysis of functional activations and have a community structure that agrees with task-based functional networks (i.e., are organized into networks such as default mode network and frontoparietal task control network). One-centimeter spherical ROIs were chosen, as they provide the largest possible size for a given ROI but preclude overlap with neighboring ROIs. Within each participant, all pairwise Pearson’s *r* correlations were calculated, yielding a 264 × 264 correlation matrix. All Pearson’s r-values were subjected to the Fisher’s z transformation to normalize the variance in correlation values. All genetic analyses used the per-participant z-correlations after regressing out several nuisance variables - scanner, gender, and summary measures of movement during the resting-state scan (average motion in the x, y, and z planes and average of roll, pitch, and yaw) for the LTS sample and gender and summary measures of movement during the resting-state scan (average motion in the x, y, and z planes and average of roll, pitch, and yaw) for the HCP sample. Gender regressor is particularly important for the HCP sample given the presence of different sex DZ twin pairs. Bivariate analyses utilized a global summary measure of each participant’s connectivity matrix which was calculated as the reciprocal of the average shortest path length between all 264 regions as calculated on a proportionally thresholded (15%) connectivity matrix using the Python package networkx [Hagberg et al., 2008]. Calculation and manipulation of connectivity matrices as well as plotting was also done in Python using the Pandas [McKinney, 2010], Seaborn (http://seaborn.pydata.org/), and Matplotlib packages [Hunter, 2007].

### Genetic Models

All genetic analyses conducted were run as structural equation models in R through the OpenMx [Boker et al., 2011] and UMX packages [Bates, 2017]. As all measures were continuous, these models utilized maximum likelihood estimation [Bentler and Weeks, 1980]. Univariate genetic models were run on each connection. A univariate model decomposes total phenotypic variation in a connection into additive genetic (A), shared environmental (C), and non-shared environmental (E) components. MZ twins share all of their genes, whereas DZ twins share on average 50% of their genes by descent, and both types are reared together. Genetic influences (A) are indicated when the MZ twin correlation is higher than the DZ correlation; shared environmental influences (C) are indicated when the DZ correlation is greater than half the MZ correlation; and non-shared environmental influences (E), which can include measurement error, are indicated when the MZ correlation is less than unity.

For the association between connections and either network connectivity, global efficiency, or movement, we utilized bivariate correlated ACE models (pictured in **Figure 6a**) and bivariate Cholesky decompositions (pictured in **Figure 6b**). The Cholesky decomposition is a common form of bivariate twin analysis, and can be used to calculate the genetic correlation and correlation predicted from A, C, and E overlap (for more see [Neale and Cardon, 1992]). In this Cholesky decomposition, the first set of A, C, and E latent variables predicting network connectivity are also allowed to predict the local connection (via paths x_2_, y_2_, and z_2_), and the local connection also has residual A, C, and E variances (obtained by squaring paths x_3_, y_3_, and z_3_). One results from this analysis is of interest. The matrix of residual A variances (i.e., squared x_3_ paths) enables us to ascertain whether the finely detailed genetic map of the connectome is simply a redescription of network connectivity (or global efficiency and movement, in those models). That is, it depicts where there are genetic influences on connections that are independent of the genetic influences on network connectivity. If residual genetic influence is present across the connectome, this analysis supports high-resolution analysis approaches as independent and complementary to analyses that utilize network-derived connectivity measures.

### Clustering Analysis

We clustered patterns of heritability estimates - rows/columns of **Figure 1a**. K-means clustering was implemented in Python using the Scikit-learn package [Pedregosa et al., 2011]. We applied clustering to the 264 × 264 matrix of A estimates to find 2-20 clusters of regions. To estimate the stability of each clustering solution, we calculated the silhouette score for each sample and averaged all scores for each clustering solution (**Figure 2a**). The silhouette score compares the distance between a region and other members of its cluster to the distance between that region and the nearest neighboring cluster in similarity space. In the current analysis, similarity was defined as the Euclidean distance between two regions’ vectors of heritability estimates. The silhouette analysis The silhouette analysis revealed several “stable” solutions in which the average silhouette score reached a local maximum, as seen in the peaks of **Figure 2a** at solutions of *k* = 2, 3, 4, 7, and 15. We describe the clustering results at the coarsest levels in the main body because they are a demonstration of novel genetic communities without the added complexity of describing many clusters.

**Figure 6.**
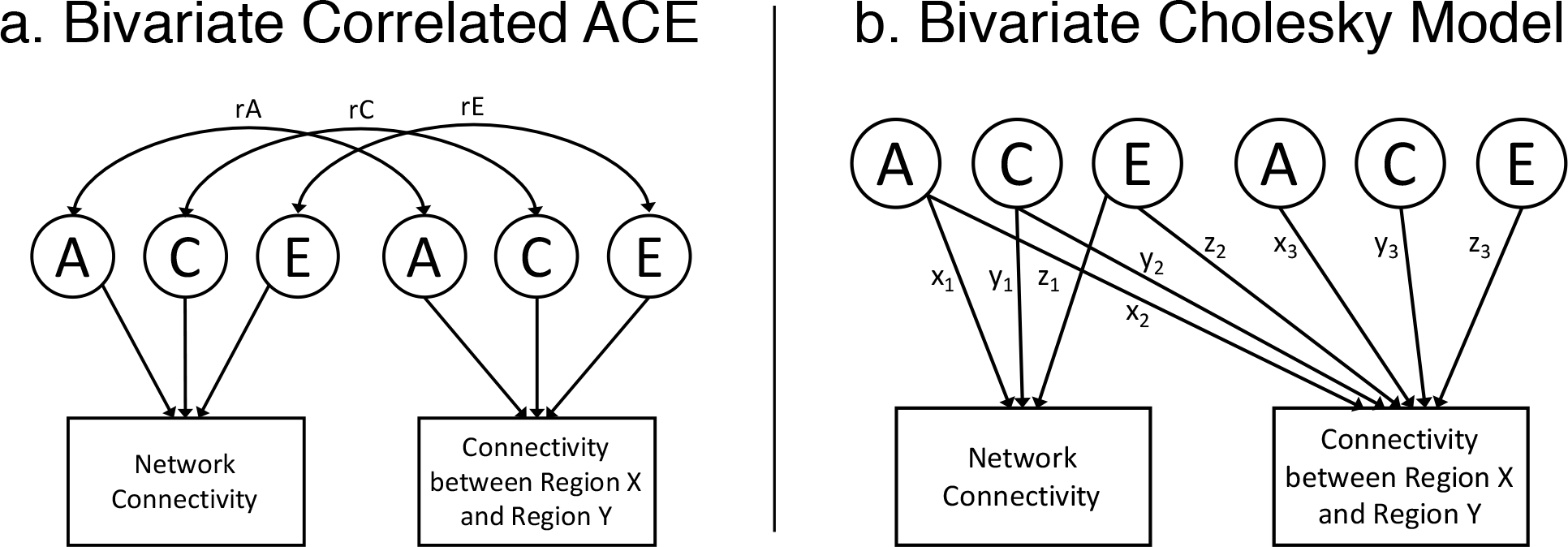
Bivariate models. a. Bivariate correlated ACE model. rA = genetic correlation. rC = shared environmental correlation. rE = non-shared environmental correlation b. Bivariate Cholesky model. Additive genetic (A), shared environmental (C), and non-shared environmental (E) latent variables (left side) predicting network connectivity (via paths x_1_, y_1_, and z_1_) and functional connectivity (via paths x_2_, y_2_, and z_2_). Functional connectivity has residual A, C, and E influences (right side). The variance explained by each influence is obtained by squaring the paths (e.g., x_3_, y_3_, and z_3_). The univariate model of network connectivity is equivalent to left side of the figure (i.e., removing the local connectivity measure).

## Data Availability

A, C, and E estimates for all pairwise connections are available for download at https://github.com/AReineberg/genetic_connectome

## Acknowledgments

This work was supported by NIH grant MH063207.

## Supporting Information

### Mean Phenotypic Connectivity

**Figures S1** and **S2** show functional connectivity averaged across all individuals in the LTS and HCP samples, respectively.

### Univatiate Twin Models - Non-shared Environmental Influence

Connection-wise estimates of non-shared environmental influence for the LTS and HCP samples are shown in **Figure S3**.

### Reliability

To examine the effect of test-retest reliability on estimation of genetic influences, we calculated edge-wise estimates of reliability utilizing multiple resting-state scans from the HCP dataset. Although a total of four resting-state scans are available for each HCP participant, we utilized only the two from the first day of scanning due to storage constraints of our computing infrastructure. Because each HCP participant’s scans were collected with different phase encoding (LR or RL with the intention of combining the two scans to minimize susceptibility artifact when compared to a single long scan using AP phase encoding, for example), we created split-scan reliability - correlation (corrected using the Spearman-Brown prophecy formula) between homologous edges from the first half of scan 1 concatenated to the second half of scan 2 and the first half of scan 2 concatenated to the second half of scan 1 - to minimize any confounding relationship between measurement error, lateralized susceptibility artifact due to LR or RL phase encoding, and start-of-scan/end-of-scan effects.

Split-scan reliability was high and normally distributed (M = 0.849, SD = 0.061). We utilized the reliability estimatess to test whether or not differences in within versus between-network connectivity persisted above and beyond any effects of reliability. We found a moderate and highly significant positive correlation between A estimates and split-scan reliability (Pearson’s *r* = 0.298, *p* < 0.001). We then regressed A estimates on within- versus between-network connectivity controlling for either reliability measure (in separate models) and found that differences in heritability estimates for within versus between-network connections persisted in both models.

### 6 vs. 30 Minutes of Resting Data

We extracted 6 minutes of resting state data for HCP participants and subjectted it the the same genetic models as we did for the full 30 minute analysis. HCP 6 minute and HCP minute A estimates were similar as quantified by correlation of one triangle of the two A estimate matrices (pearsonr = 0.349, p = 0.0). HCP 6 minute and LTS estimates were less similar (Pearson’s r = 0.087, p = 5.57e-59)

### Within- versus Between-network Connectiity

We quantified differences in heritability for within- and between-network connections summazired at various levels of analysis - averaged across the whole connectome (**Table S1a**), averaged for within- and between-network connections for 14 *a priori* networks of interest (**Table S1b**), and for each of 264 regions that are part of the parcellation utilized in the current study (**Table S2**). These averages included all estimates, even those estimated as zero.

### Clustering

**Figure S5** shows the configuration of the *k* = 15 clustering solution for the LTS sample. Although this analysis is not heirarchically related to the *k* = 3 solution in a quantitative sense, the three super clusters described earlier qualitatively break down into smaller pieces in this analysis (referred to as subclusters here). Each supercluster maintains a large subcluster in the *k* = 15 solution (clusters 1, 2, and 3 correspond to superslusters 1, 2, and 3). There are 12 subclusters which are stable, smaller subcomponents of the superclusters - subclusters 4 through 9 correspond to supercluster 1; subcluster 10 corresponds to supercluster 2; and subclusters 11 through 15 correspond to supercluster 3. Again, because the k-means algorithm is not heirarchical in nature, the mapping of subclusters to subclusters is not perfect. In summary, this analysis reveals 1. a correspondence between genetic communities and classic resting-state functional communities (e.g., distinct grouping of sensory versus cognitive regions) and 2. a genetic taxonomy of functional brain regions that is novel as evidenced by small subclusters with regions of diverse function but similar patterns of heritable connectivity.

We subjected the HCP sample heritability matrix to the same clustering procedure used with the LTS sample. We found stable solutions at *k*-values of 2, 3, 7, and 13 (**Figure S6a**). For the most direct comparison to the LTS *k* = 3 supercluster solution, we investigated the *k* = 3 solution in the HCP sample in great detail. The *k* = 3 clustering solution is shown in **Figure S6b**. The super-clusters of the HCP sample were similar in nature to those of the LTS sample, with one super-cluster containing regions with no particualrly clear pattern of heritable connectivity (1), another supercluster containing regions with particularly heritable connectivity to sensory areas (2; predominantly visual regions), and a third super-cluster with particularly heritable connectivity to default and other cognitive regions (3). **Figure S7** shows the spatial locations of and *a priori* network assignments for regions that are part of the three LTS super-clusters.

### Bivariate Analyses

To ascertain whether global efficiency captures most of the genetic variance in local connectivity (*question 3*), we conducted a bivariate genetic analysis of local connections and global efficiency. These bivariate analyses quantify the degree to which local functional connectivity is genetically separable from a summary measure of the connectome. We found global efficiency was more heritable in the HCP sample than in the LTS sample. In the HCP sample, 36.6% of the variance in global efficiency was attributed to additive genetic influences. In the LTS sample, 6.3% of the variance in global efficiency was attributed to additive genetic influences. Given this difference, estimates of global efficiency may be sensitive to the short scan length or the smaller sample size of the LTS data.

Residual additive genetic influence after accounting for genetic influence shared with global efficiency (i.e., residual A variances or specific A) for each functional connection (i.e., squared x_3_ paths; see **Figure 6**) was distributed bi-modally in a manner similar to the univariate A estimates. In the HCP sample, approximately 64.5% of connections (22,400) had zero residual A variance with a separate positively skewed distribution describing the remaining 12,316 connections (M = 0.117, SD = 0.073, Skew = 0.726, Range = 0.010 - 0.524). In the LTS sample, approximately 75.8% of connections (26,318) had zero residual A variance, with a separate positively skewed distribution describing the remaining 8,398 connections (M = 0.127, SD = 0.087, Skew = 0.926, Range = 0.010 - 0.520). The matrix of specific A estimates is displayed in **Figure S8** for the HCP and LTS samples (lower and upper triangle respectively). These results suggest that there are genetic influences specific to local functional connections. That is, the genetic influences on finer-grained measures of functional connections are not simply reflecting the genetic influences on global efficiency. Future work should synthesize the spatial distribution of these global and specific effects.

**Figure S1.**
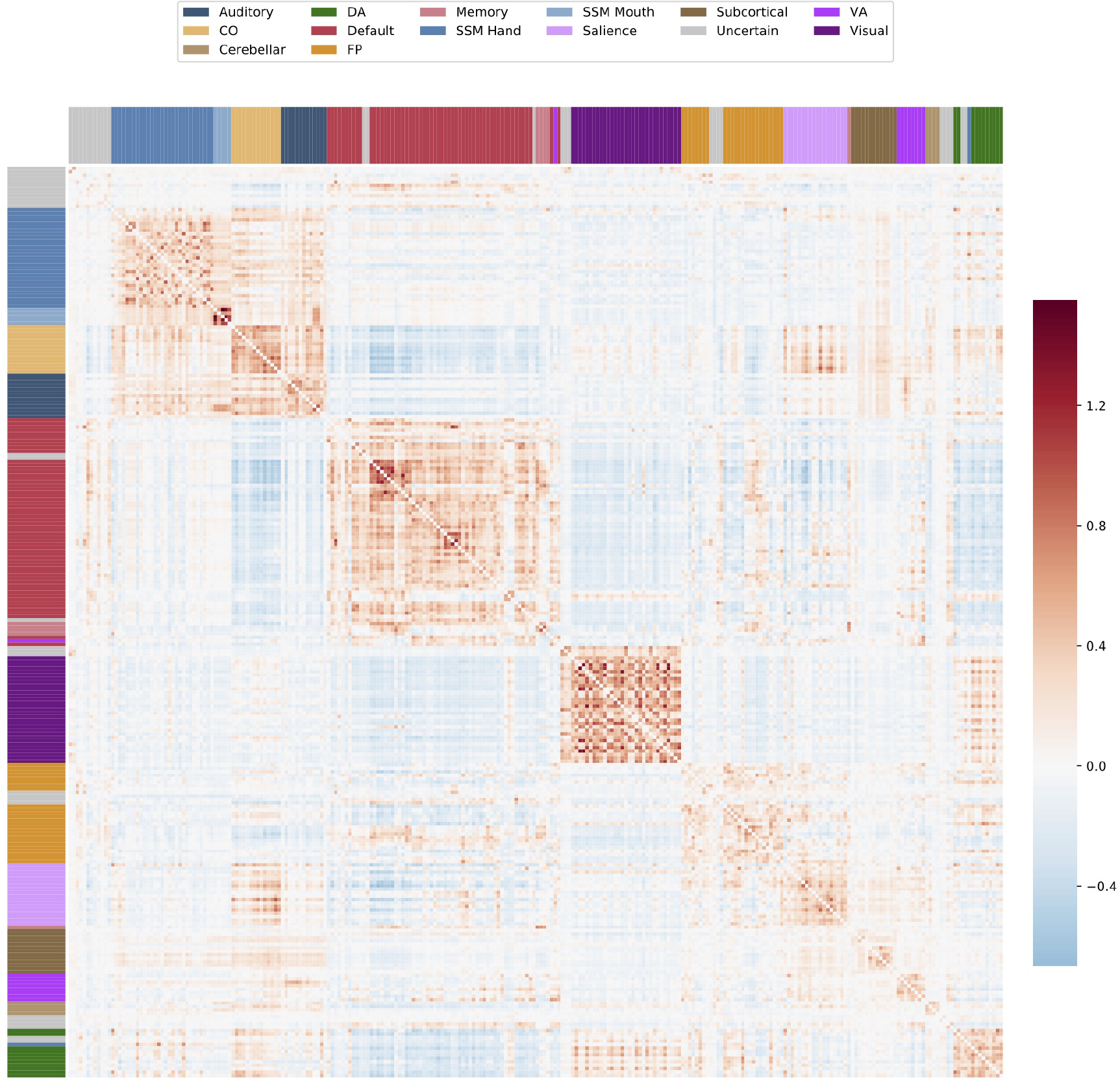
Mean phenotypic connectivity across LTS participants. Group average connectome for LTS sample. CO = cingulo-opercular, DA = dorsal attention, FP = frontoparietal, SSM = sensory/somatomotor, VA = ventral attention.

**Figure S2.**
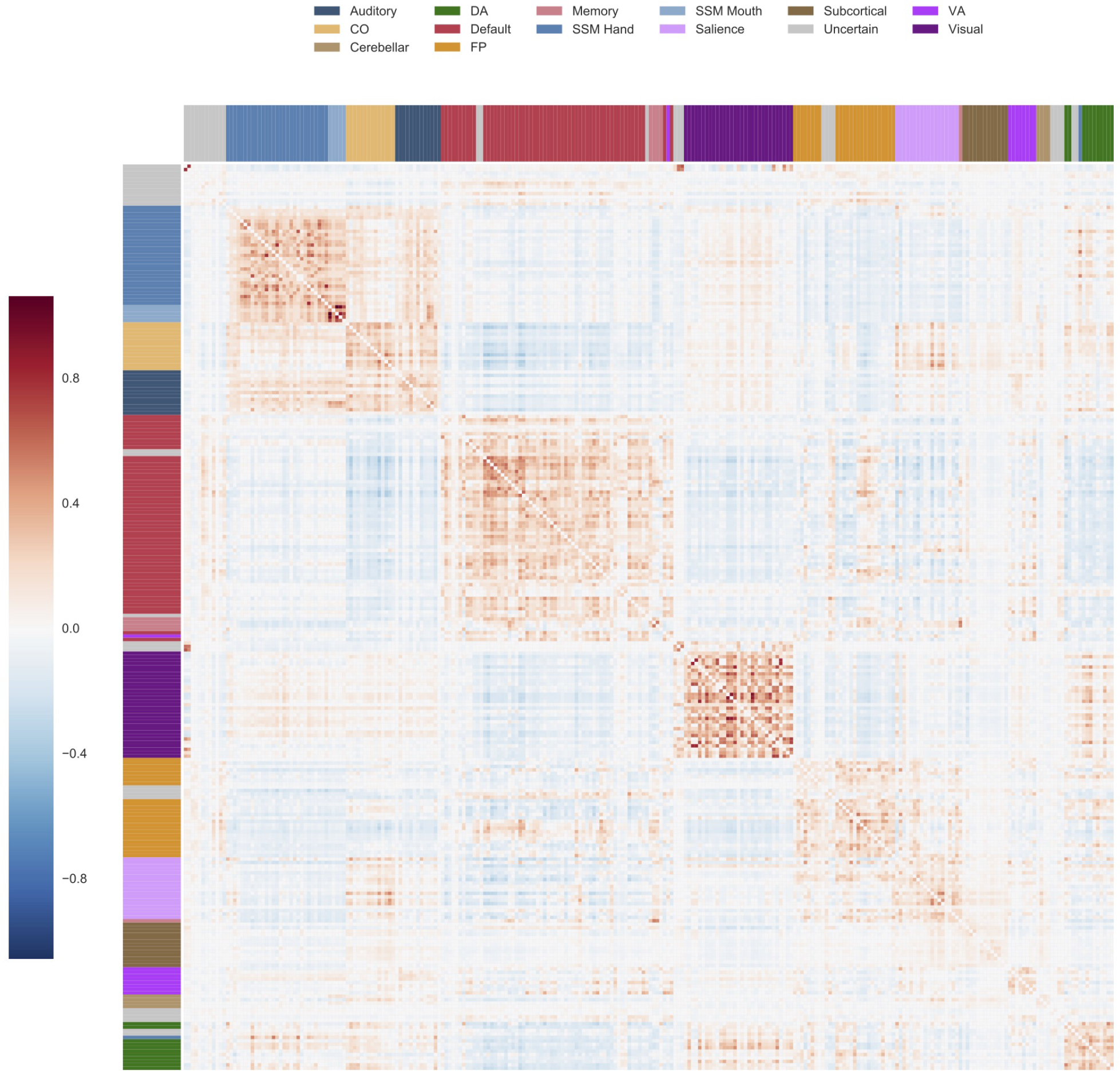
Mean phenotypic connectivity across HCP participants. Group average connectome for HCP sample. CO = cingulo-opercular, DA = dorsal attention, FP = frontoparietal, SSM = sensory/somatomotor, VA = ventral attention.

**Figure S3.**
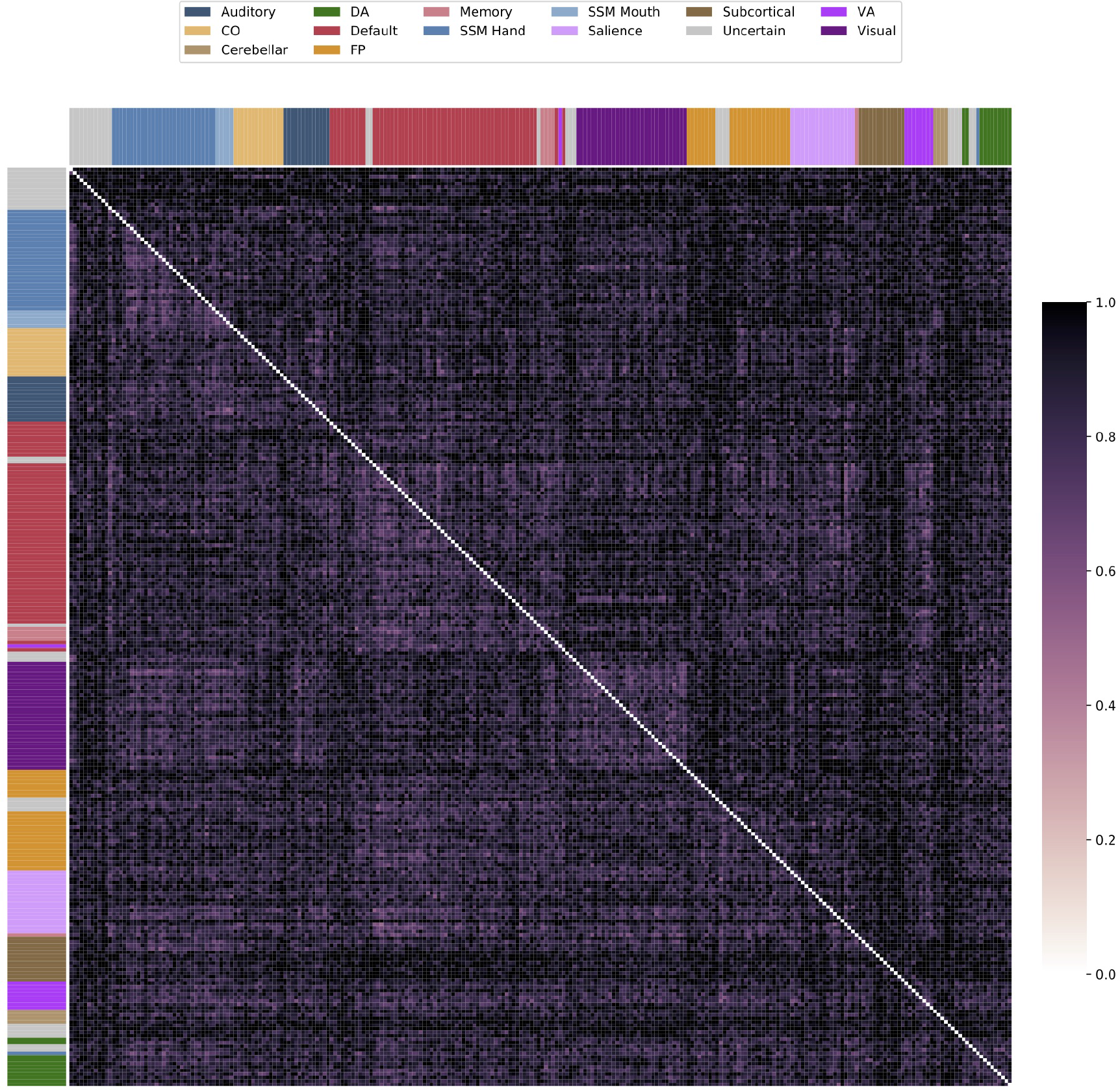
Connection-wise estimates of non-shared environmental influence in LTS and HCP samples. Estimates of nonshared environmental influence (E) are displayed for the HCP and LTS samples in the lower and upper triangles, respectively. CO = cingulo-opercular, DA = dorsal attention, FP = frontoparietal, SSM = sensory/somatomotor, VA = ventral attention.

**Figure S4.**
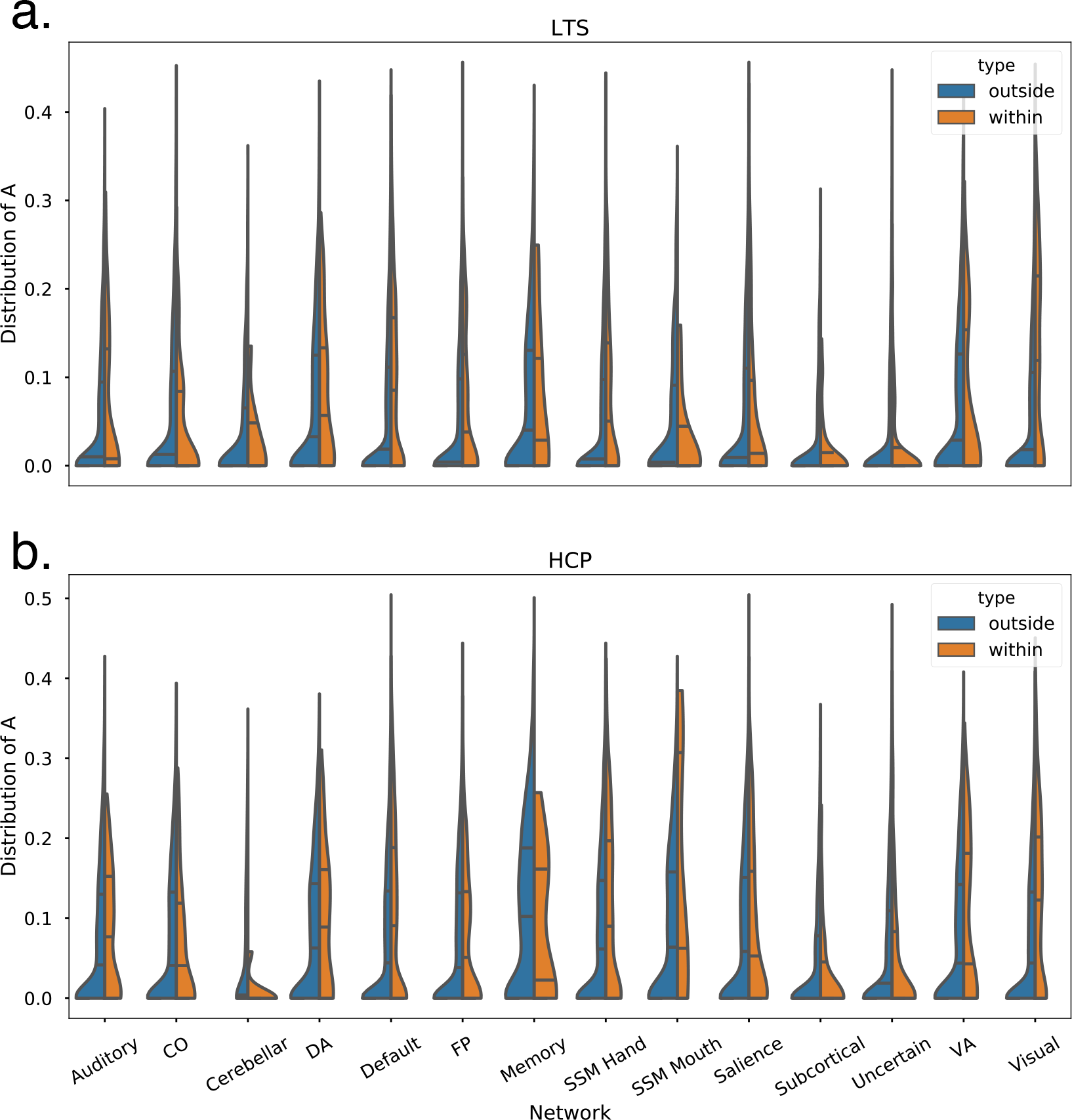
Connection-wise estimates of additive genetic influence summarized by *a priori* networks. Violin plots of distribution of additive genetic (A) estimates grouped by *a priori* networks reveal higher heritability for within- than between-network connections in both LTS (a) and HCP (b) samples. CO = cingulo-opercular, DA = dorsal attention, FP = frontoparietal, SSM = sensory/somatomotor, VA = ventral attention.

**Table 1.** Within- versus between network connection heritability by network

|  | LTS |  |  |  | HCP |  |  |  |
| --- | --- | --- | --- | --- | --- | --- | --- | --- |
|  | m<sub>within</sub> | m<sub>between</sub> | *t* | *p* | m<sub>within</sub> | m<sub>between</sub> | *t* | *p* |
| a. Heritability (A)<sub>Whole Connectome</sub> |  |  |  |  |  |  |  |  |
| - | 0.088 | 0.056 | 19.282 | <0.001 | 0.101 | 0.075 | 15.111 | <0.001 |
| b. Heritability (A) by network |  |  |  |  |  |  |  |  |
| Memory | 0.094 | 0.072 | 0.757 | ns | 0.107 | 0.112 | -0.160 | ns |
| FP | 0.072 | 0.054 | 3.717 | 0.000 | 0.079 | 0.072 | 1.536 | ns |
| SSM Mouth | 0.041 | 0.053 | -0.645 | ns | 0.171 | 0.088 | 1.726 | ns |
| Salience | 0.065 | 0.062 | 0.524 | ns | 0.092 | 0.085 | 0.821 | ns |
| DA | 0.089 | 0.069 | 1.699 | 0.095 | 0.110 | 0.082 | 2.176 | 0.034 |
| Visual | 0.131 | 0.060 | 13.732 | 0.000 | 0.126 | 0.074 | 10.533 | 0.000 |
| VA | 0.078 | 0.071 | 0.483 | ns | 0.110 | 0.077 | 1.828 | 0.076 |
| Auditory | 0.074 | 0.055 | 1.912 | 0.060 | 0.089 | 0.073 | 1.760 | 0.082 |
| Default | 0.101 | 0.063 | 15.428 | 0.000 | 0.110 | 0.076 | 12.712 | 0.000 |
| Cerebellar | 0.039 | 0.040 | -0.021 | ns | 0.010 | 0.053 | -4.356 | 0.006 |
| SSM Hand | 0.084 | 0.054 | 6.782 | 0.000 | 0.116 | 0.085 | 5.833 | 0.000 |
| Subcortical | 0.020 | 0.032 | -2.728 | 0.008 | 0.035 | 0.047 | -1.788 | 0.078 |
| Uncertain | 0.025 | 0.039 | -5.461 | 0.000 | 0.053 | 0.062 | -2.228 | 0.026 |
| CO | 0.051 | 0.061 | -1.408 | ns | 0.073 | 0.073 | 0.046 | ns |
| c. Shared-environmental influence (C) by network |  |  |  |  |  |  |  |  |
| Memory | 0.078 | 0.038 | 1.237 | ns | 0.063 | 0.039 | 0.934 | ns |
| FP | 0.036 | 0.032 | 1.188 | ns | 0.045 | 0.040 | 1.324 | ns |
| SSM Mouth | 0.136 | 0.026 | 4.527 | 0.001 | 0.112 | 0.046 | 2.156 | 0.059 |
| Salience | 0.040 | 0.037 | 0.608 | ns | 0.028 | 0.037 | -2.165 | 0.032 |
| DA | 0.039 | 0.032 | 0.738 | ns | 0.040 | 0.035 | 0.571 | ns |
| Visual | 0.054 | 0.033 | 6.410 | 0.000 | 0.047 | 0.040 | 2.278 | 0.023 |
| VA | 0.060 | 0.044 | 1.189 | ns | 0.029 | 0.048 | -1.839 | 0.074 |
| Auditory | 0.044 | 0.033 | 1.476 | ns | 0.038 | 0.041 | -0.470 | ns |
| Default | 0.030 | 0.035 | -2.938 | 0.003 | 0.049 | 0.037 | 6.786 | 0.000 |
| Cerebellar | 0.042 | 0.022 | 0.792 | ns | 0.036 | 0.028 | 0.360 | ns |
| SSM Hand | 0.026 | 0.030 | -1.700 | 0.090 | 0.055 | 0.038 | 4.438 | 0.000 |
| Subcortical | 0.033 | 0.026 | 1.262 | ns | 0.041 | 0.032 | 1.348 | ns |
| Uncertain | 0.020 | 0.026 | -2.516 | 0.012 | 0.027 | 0.034 | -2.737 | 0.006 |
| CO | 0.046 | 0.036 | 1.696 | 0.093 | 0.052 | 0.039 | 1.706 | 0.091 |

**Table 2.**
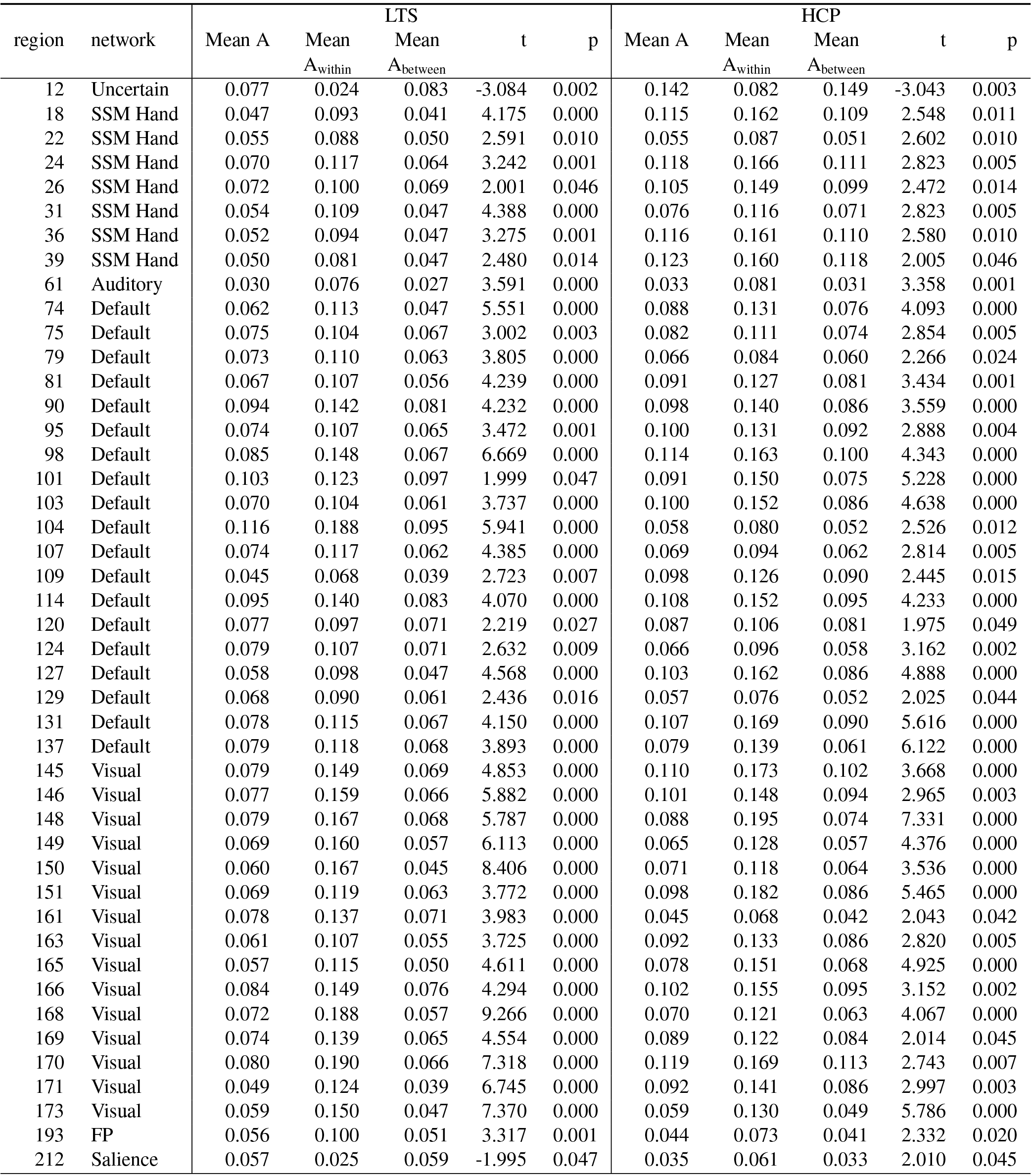
Within- versus between-network connection heritability by region. Region is 0-indexed

| region | network | LTS |  |  |  |  | HCP |  |  |  |  |
| --- | --- | --- | --- | --- | --- | --- | --- | --- | --- | --- | --- |
|  |  | Mean A | Mean<br>A<sub>within</sub> | Mean<br>A<sub>between</sub> | t | p | Mean A | Mean<br>A<sub>within</sub> | Mean<br>A<sub>between</sub> | t | p |
| 12 | Uncertain | 0.077 | 0.024 | 0.083 | -3.084 | 0.002 | 0.142 | 0.082 | 0.149 | -3.043 | 0.003 |
| 18 | SSM Hand | 0.047 | 0.093 | 0.041 | 4.175 | 0.000 | 0.115 | 0.162 | 0.109 | 2.548 | 0.011 |
| 22 | SSM Hand | 0.055 | 0.088 | 0.050 | 2.591 | 0.010 | 0.055 | 0.087 | 0.051 | 2.602 | 0.010 |
| 24 | SSM Hand | 0.070 | 0.117 | 0.064 | 3.242 | 0.001 | 0.118 | 0.166 | 0.111 | 2.823 | 0.005 |
| 26 | SSM Hand | 0.072 | 0.100 | 0.069 | 2.001 | 0.046 | 0.105 | 0.149 | 0.099 | 2.472 | 0.014 |
| 31 | SSM Hand | 0.054 | 0.109 | 0.047 | 4.388 | 0.000 | 0.076 | 0.116 | 0.071 | 2.823 | 0.005 |
| 36 | SSM Hand | 0.052 | 0.094 | 0.047 | 3.275 | 0.001 | 0.116 | 0.161 | 0.110 | 2.580 | 0.010 |
| 39 | SSM Hand | 0.050 | 0.081 | 0.047 | 2.480 | 0.014 | 0.123 | 0.160 | 0.118 | 2.005 | 0.046 |
| 61 | Auditory | 0.030 | 0.076 | 0.027 | 3.591 | 0.000 | 0.033 | 0.081 | 0.031 | 3.358 | 0.001 |
| 74 | Default | 0.062 | 0.113 | 0.047 | 5.551 | 0.000 | 0.088 | 0.131 | 0.076 | 4.093 | 0.000 |
| 75 | Default | 0.075 | 0.104 | 0.067 | 3.002 | 0.003 | 0.082 | 0.111 | 0.074 | 2.854 | 0.005 |
| 79 | Default | 0.073 | 0.110 | 0.063 | 3.805 | 0.000 | 0.066 | 0.084 | 0.060 | 2.266 | 0.024 |
| 81 | Default | 0.067 | 0.107 | 0.056 | 4.239 | 0.000 | 0.091 | 0.127 | 0.081 | 3.434 | 0.001 |
| 90 | Default | 0.094 | 0.142 | 0.081 | 4.232 | 0.000 | 0.098 | 0.140 | 0.086 | 3.559 | 0.000 |
| 95 | Default | 0.074 | 0.107 | 0.065 | 3.472 | 0.001 | 0.100 | 0.131 | 0.092 | 2.888 | 0.004 |
| 98 | Default | 0.085 | 0.148 | 0.067 | 6.669 | 0.000 | 0.114 | 0.163 | 0.100 | 4.343 | 0.000 |
| 101 | Default | 0.103 | 0.123 | 0.097 | 1.999 | 0.047 | 0.091 | 0.150 | 0.075 | 5.228 | 0.000 |
| 103 | Default | 0.070 | 0.104 | 0.061 | 3.737 | 0.000 | 0.100 | 0.152 | 0.086 | 4.638 | 0.000 |
| 104 | Default | 0.116 | 0.188 | 0.095 | 5.941 | 0.000 | 0.058 | 0.080 | 0.052 | 2.526 | 0.012 |
| 107 | Default | 0.074 | 0.117 | 0.062 | 4.385 | 0.000 | 0.069 | 0.094 | 0.062 | 2.814 | 0.005 |
| 109 | Default | 0.045 | 0.068 | 0.039 | 2.723 | 0.007 | 0.098 | 0.126 | 0.090 | 2.445 | 0.015 |
| 114 | Default | 0.095 | 0.140 | 0.083 | 4.070 | 0.000 | 0.108 | 0.152 | 0.095 | 4.233 | 0.000 |
| 120 | Default | 0.077 | 0.097 | 0.071 | 2.219 | 0.027 | 0.087 | 0.106 | 0.081 | 1.975 | 0.049 |
| 124 | Default | 0.079 | 0.107 | 0.071 | 2.632 | 0.009 | 0.066 | 0.096 | 0.058 | 3.162 | 0.002 |
| 127 | Default | 0.058 | 0.098 | 0.047 | 4.568 | 0.000 | 0.103 | 0.162 | 0.086 | 4.888 | 0.000 |
| 129 | Default | 0.068 | 0.090 | 0.061 | 2.436 | 0.016 | 0.057 | 0.076 | 0.052 | 2.025 | 0.044 |
| 131 | Default | 0.078 | 0.115 | 0.067 | 4.150 | 0.000 | 0.107 | 0.169 | 0.090 | 5.616 | 0.000 |
| 137 | Default | 0.079 | 0.118 | 0.068 | 3.893 | 0.000 | 0.079 | 0.139 | 0.061 | 6.122 | 0.000 |
| 145 | Visual | 0.079 | 0.149 | 0.069 | 4.853 | 0.000 | 0.110 | 0.173 | 0.102 | 3.668 | 0.000 |
| 146 | Visual | 0.077 | 0.159 | 0.066 | 5.882 | 0.000 | 0.101 | 0.148 | 0.094 | 2.965 | 0.003 |
| 148 | Visual | 0.079 | 0.167 | 0.068 | 5.787 | 0.000 | 0.088 | 0.195 | 0.074 | 7.331 | 0.000 |
| 149 | Visual | 0.069 | 0.160 | 0.057 | 6.113 | 0.000 | 0.065 | 0.128 | 0.057 | 4.376 | 0.000 |
| 150 | Visual | 0.060 | 0.167 | 0.045 | 8.406 | 0.000 | 0.071 | 0.118 | 0.064 | 3.536 | 0.000 |
| 151 | Visual | 0.069 | 0.119 | 0.063 | 3.772 | 0.000 | 0.098 | 0.182 | 0.086 | 5.465 | 0.000 |
| 161 | Visual | 0.078 | 0.137 | 0.071 | 3.983 | 0.000 | 0.045 | 0.068 | 0.042 | 2.043 | 0.042 |
| 163 | Visual | 0.061 | 0.107 | 0.055 | 3.725 | 0.000 | 0.092 | 0.133 | 0.086 | 2.820 | 0.005 |
| 165 | Visual | 0.057 | 0.115 | 0.050 | 4.611 | 0.000 | 0.078 | 0.151 | 0.068 | 4.925 | 0.000 |
| 166 | Visual | 0.084 | 0.149 | 0.076 | 4.294 | 0.000 | 0.102 | 0.155 | 0.095 | 3.152 | 0.002 |
| 168 | Visual | 0.072 | 0.188 | 0.057 | 9.266 | 0.000 | 0.070 | 0.121 | 0.063 | 4.067 | 0.000 |
| 169 | Visual | 0.074 | 0.139 | 0.065 | 4.554 | 0.000 | 0.089 | 0.122 | 0.084 | 2.014 | 0.045 |
| 170 | Visual | 0.080 | 0.190 | 0.066 | 7.318 | 0.000 | 0.119 | 0.169 | 0.113 | 2.743 | 0.007 |
| 171 | Visual | 0.049 | 0.124 | 0.039 | 6.745 | 0.000 | 0.092 | 0.141 | 0.086 | 2.997 | 0.003 |
| 173 | Visual | 0.059 | 0.150 | 0.047 | 7.370 | 0.000 | 0.059 | 0.130 | 0.049 | 5.786 | 0.000 |
| 193 | FP | 0.056 | 0.100 | 0.051 | 3.317 | 0.001 | 0.044 | 0.073 | 0.041 | 2.332 | 0.020 |
| 212 | Salience | 0.057 | 0.025 | 0.059 | -1.995 | 0.047 | 0.035 | 0.061 | 0.033 | 2.010 | 0.045 |

**Figure S5.**
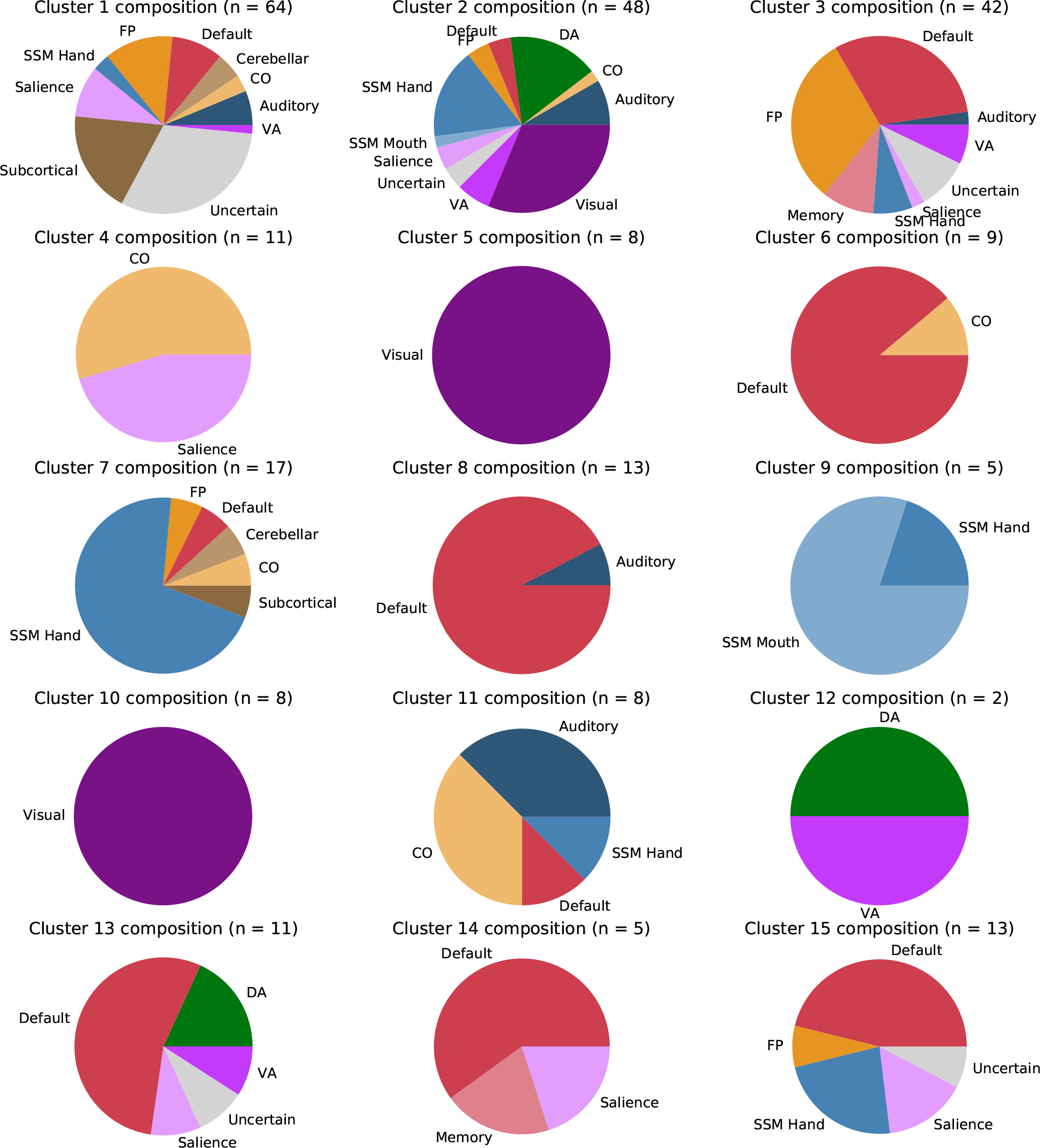
K-means 15-cluster configurations. Cluster configurations from LTS sample *k* = 15 clustering solution. n = number of regions, CO = cingulo-opercular, DA = dorsal attention, FP = frontoparietal, SSM = sensory/somatomotor, VA ventral attention.

**Figure S6.**
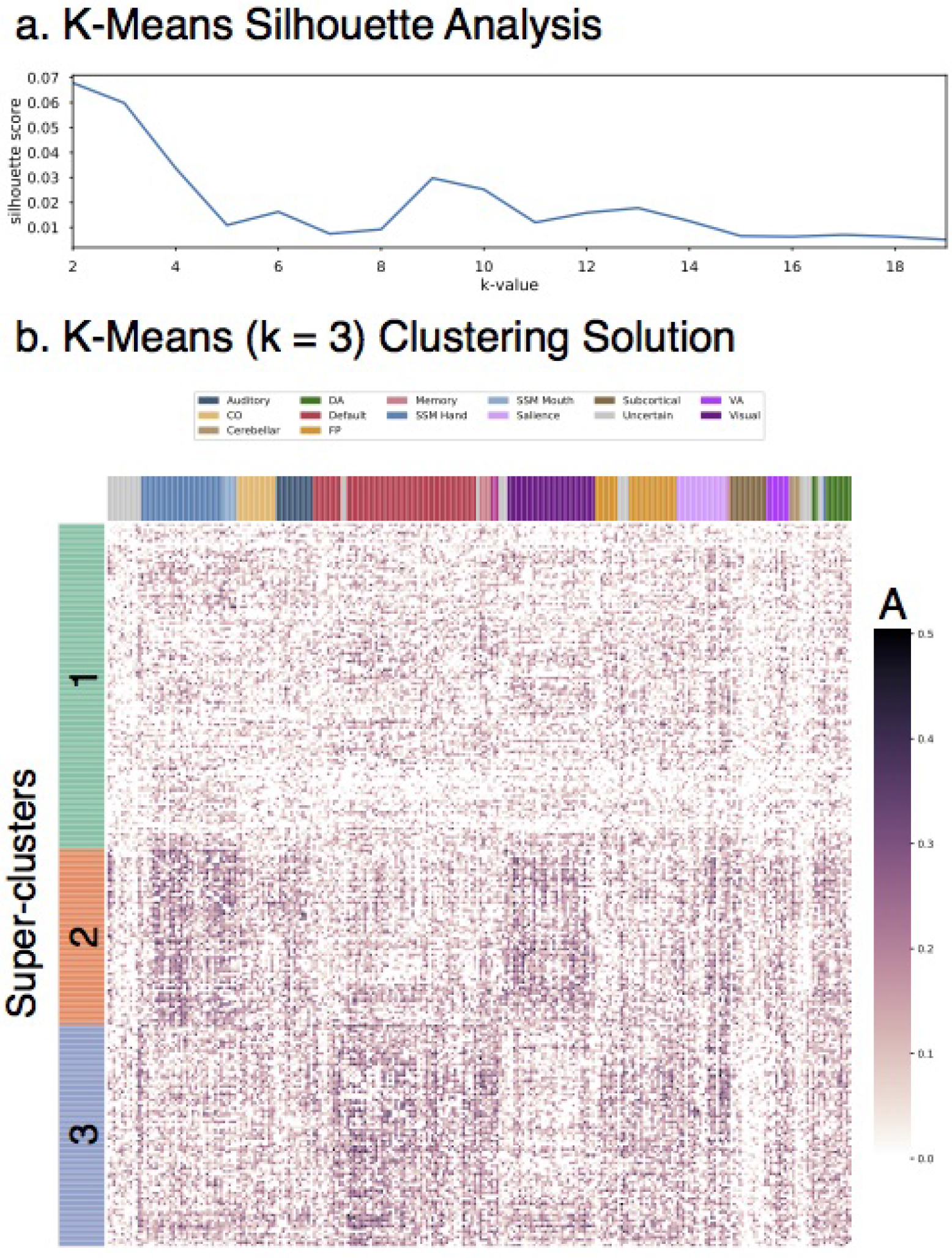
K-means 3-cluster solution (HCP sample) Row-wise clustering of HCP additive genetic (A) estimates reveals several stable clustering solutions of regions with similar patterns of connectivity heritability. Super-clusters (*k* = 3) are described in detail for comparison to resuls in the LTS sample. a. Silhouette analysis revels stable clustering solutions at k-values of 2, 3, 7, and 13. b. Clustered version of HCP A estimate matrix for *k* = 3 solution. CO = cingulo-opercular, DA = dorsal attention, FP = frontoparietal, SSM = sensory/somatomotor, VA = ventral attention.

**Figure S7.**
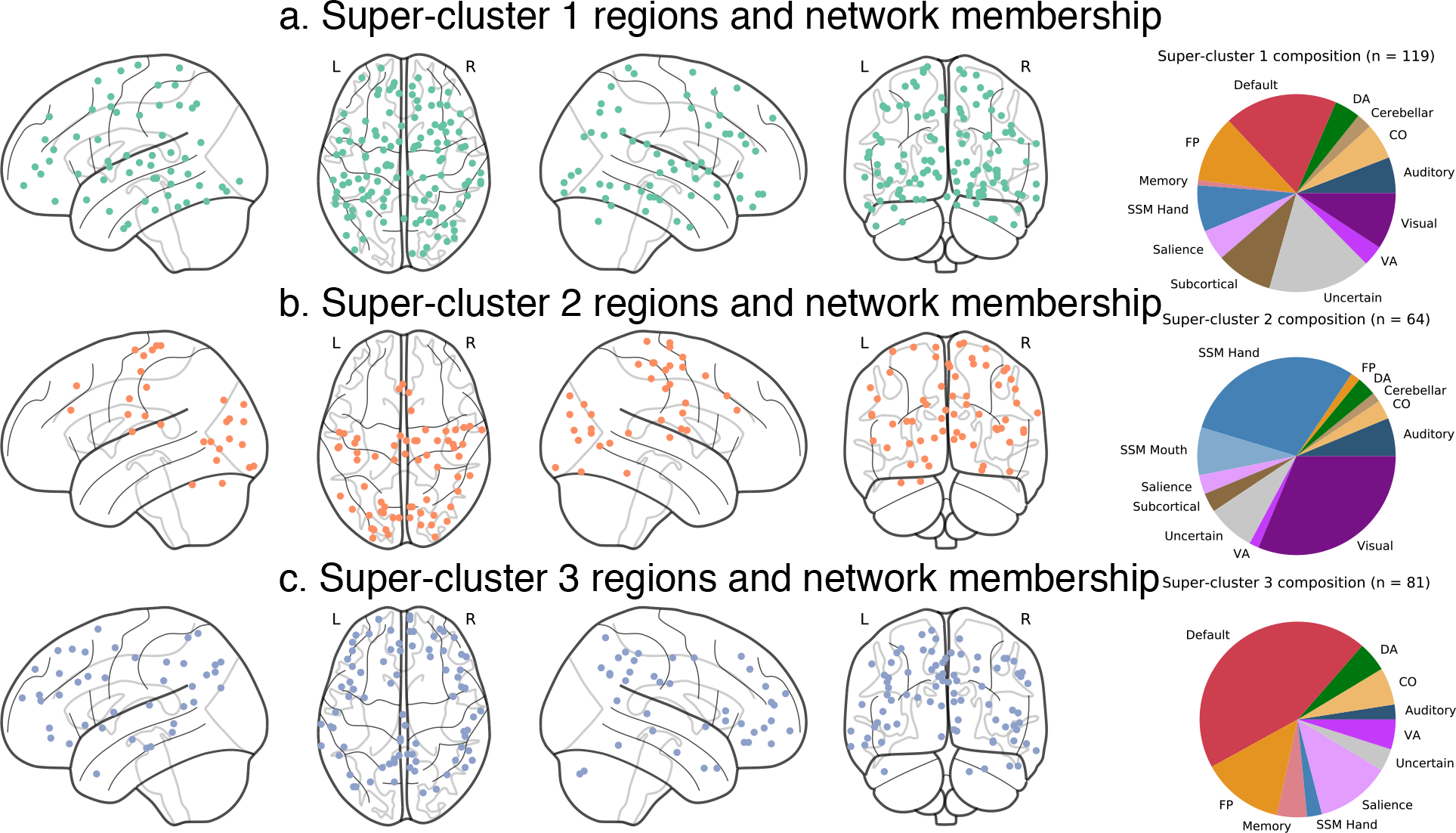
K-means 3-cluster summary (HCP sample) Spatial location of regions from super-clusters 1-3 of *k* = 3 solution in the LTS sample. a. Super-cluster 1 regions are widely distributed across the brain. b. Super-cluster 2 regions are located across lateral prefrontal, lateral parietal, mid and anterior temporal, midline frontal, and cingulate areas. c. Super-cluster 3 regions are located primarily in visual areas. n = number of regions, CO = cingulo-opercular, DA = dorsal attention, FP = frontoparietal, SSM = sensory/somatomotor, VA = ventral attention.

**Figure S8.**
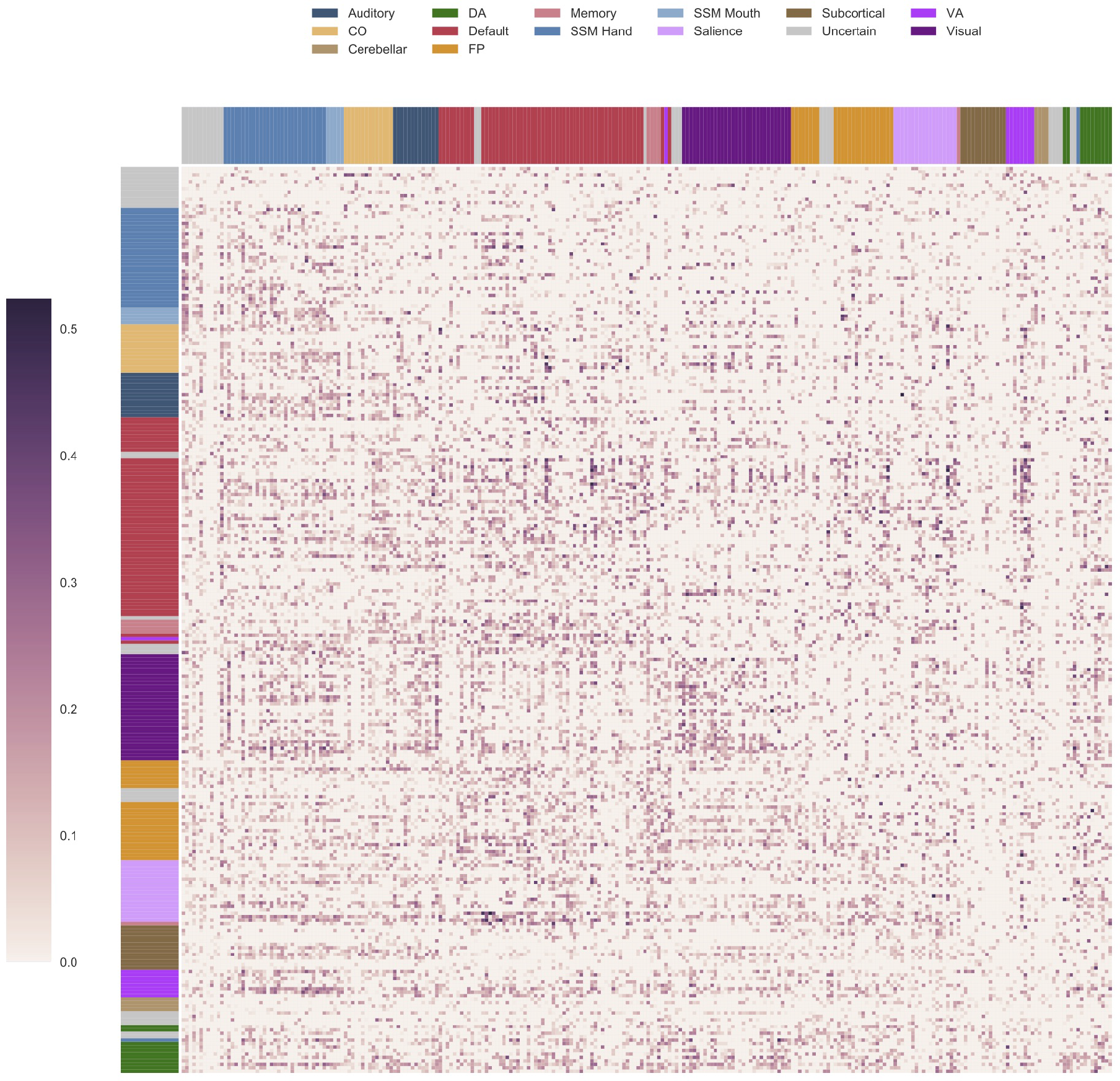
Genetic Variance in Connections After Accounting for Global Efficiency. Estimates of residual genetic variance for each connection after accounting for genetic influence shared with global efficiency. Estimates for HCP and LTS samples are in lower and upper triangles, respectively. CO = cingulo-opercular, DA = dorsal attention, FP = frontoparietal, SSM = sensory/somatomotor, VA = ventral attention.

**Figure S9.**
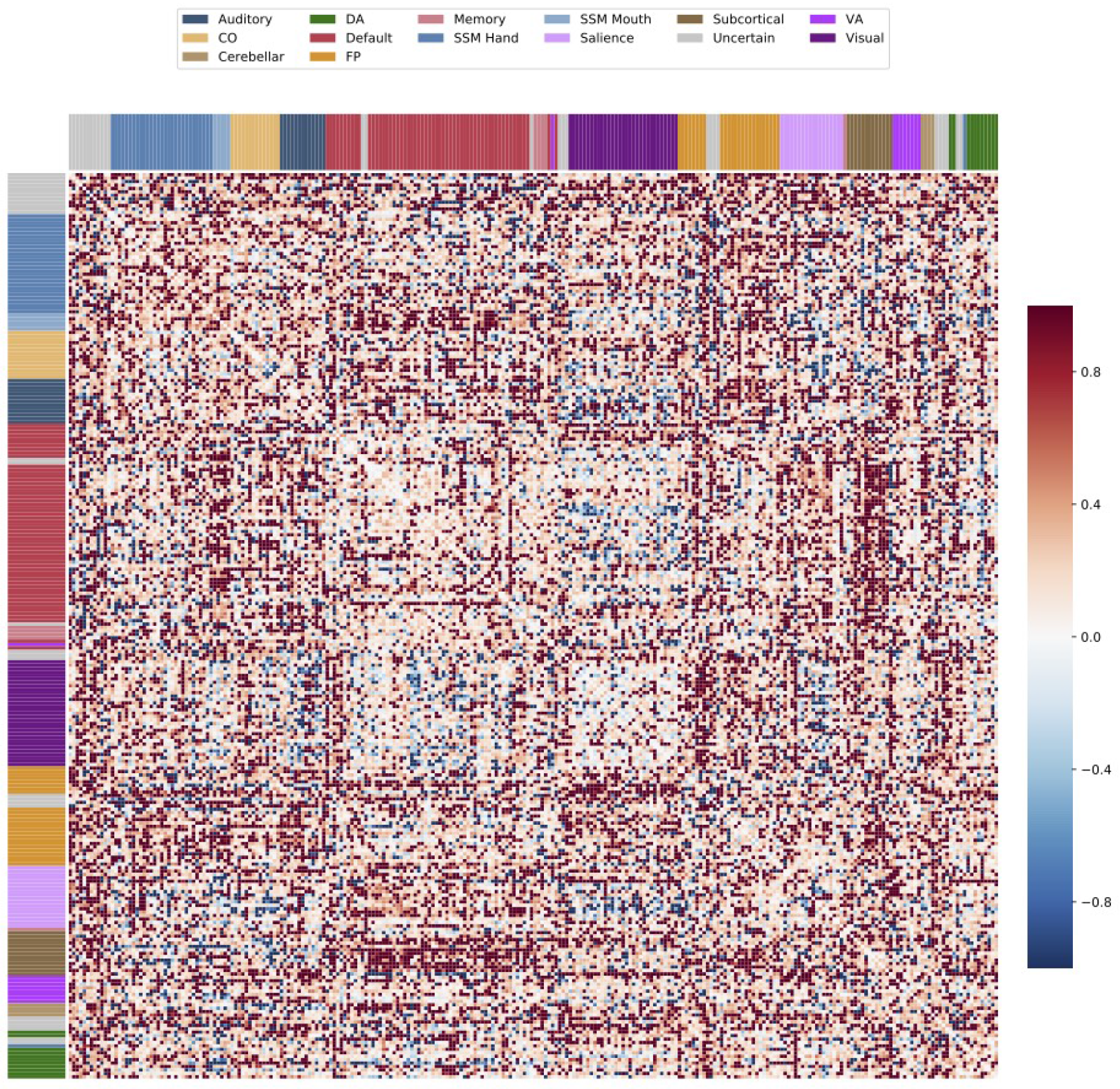
Genetic Correlation of Connections and In-Scanner Movement. Each cell of the matrix represents the genetic correlation between a connection and rotation movement.

